# Single-cell RNA-sequencing of peripheral neuroblastic tumors reveals an aggressive transitional cell state at the junction of an adrenergic-mesenchymal transdifferentiation trajectory

**DOI:** 10.1101/2020.05.15.097469

**Authors:** Xiaojun Yuan, Janith A. Seneviratne, Shibei Du, Ying Xu, Yijun Chen, Qianya Jin, Xuanxuan Jin, Anushree Balachandran, Shihao Huang, Yanli Xu, Yue Zhai, Liumei Lu, Mengjie Tang, Yushuang Dong, Belamy B. Cheung, Glenn M. Marshall, Weiyang Shi, Chao Zhang, Daniel R. Carter

## Abstract

Peripheral neuroblastic tumors (PNTs) are the most common extracranial solid tumors in early childhood. They represent a spectrum of neural crest derived tumors including neuroblastoma, ganglioneuroblastoma and ganglioneuroma. PNTs exhibit heterogeneity due to interconverting malignant cell states described as adrenergic/nor-adrenergic or mesenchymal/neural crest cell in origin. The factors determining individual patient levels of tumor heterogeneity, their impact on the malignant phenotype, and the presence of other cell states are unknown. Here, single-cell RNA-sequencing analysis of 4267 cells from 7 PNTs demonstrated extensive transcriptomic heterogeneity. Trajectory modelling showed that malignant neuroblasts move between adrenergic and mesenchymal cell states via a novel state that we termed a “transitional” phenotype. Transitional cells are characterized by gene expression programs linked to a sympathoadrenal development, and aggressive tumor phenotypes such as rapid proliferation and tumor dissemination. Among primary bulk tumor patient cohorts, high expression of the transitional gene signature was highly predictive of poor prognosis when compared to adrenergic and mesenchymal expression patterns. High transitional gene expression in neuroblastoma cell lines identified a similar transitional H3K27-acetylation super-enhancer landscape, supporting the concept that PNTs have phenotypic plasticity and transdifferentiation capacity. Additionally, examination of PNT microenvironments, found that neuroblastomas contained low immune cell infiltration, high levels of non-inflammatory macrophages, and low cytotoxic T lymphocyte levels compared with more benign PNT subtypes. Modeling of cell-cell signaling in the tumor microenvironment predicted specific paracrine effects toward the various subtypes of malignant cells, suggesting further cell-extrinsic influences on malignant cell phenotype. Collectively, our study reveals the presence of a previously unrecognized transitional cell state with high malignant potential and an immune cell architecture which serve both as potential biomarkers and therapeutic targets.

## Introduction

Peripheral neuroblastic tumors (PNTs) represent a spectrum of tumors derived from the neural crest and account up to 8-10% of all pediatric malignancies. A salient feature of PNTs is a heterogeneous clinical course ranging from spontaneous regression to persistent disease progression^1^. Histologically, PNTs comprise four variants, including neuroblastoma (NB), ganglioneuroblastoma nodular (GNBn), ganglioneuroblastoma intermixed (GNBi), and ganglioneuroma (GN)^1^. GN and GNBi are low-grade in nature and usually curable by surgical resection alone^2^. In contrast, the most common subtype; NB is often lethal. Despite intensive treatments, the long-term survival of high-risk NB is less than 50%^3^. Around half of high-risk patients relapse after initial treatment response, and salvage therapies for relapsed patients are rarely effective^1^. Moreover, genomics studies comparing longitudinal samples from the same patient show that clonal evolution is prominent feature of disease progression^4-6^. Therefore, a better understanding of tumor heterogeneity will be required to improve therapy for patients.

Emerging evidence proposes that tumor plasticity in neuroblastic cells contributes to chemoresistance^7,8^. These studies conjointly suggest that neuroblastoma is composed of transdifferentiating malignant cells with distinct epigenetic and transcriptomic landscapes. In these studies, tumors and neuroblastoma cell lines were subtyped according to a two-group classification conforming to an adrenergic/noradrenergic state or a mesenchymal/neural crest cell state. Importantly, cell state was seen to be relevant for therapeutic efficacy, with mesenchymal neuroblastic cells more resistant to conventional anti-cancer therapies^8^. Nevertheless, while these studies demonstrate the importance of cell plasticity in neuroblastoma, bulk-tissue profiling is still compromised by averaging the individual impact of cell types within the tumor microenvironment, potentially hindering the identification of other cell types.

To assess intratumoral compositions across PNT subtypes, we conducted single-cell RNA sequencing (scRNA-seq) on more than 4000 cells of fresh, surgically resected tissues from 7 PNT patients, which spanned all histological variants, including 3 NB, 2 GNBn, 1 GNBi and 1 GN lesion. Our results suggest that malignant neuroblasts can exist as an intermediate between adrenergic and mesenchymal transcriptional states via a previously unidentified transitional population. Transitional neuroblasts have an aggressive neurodevelopmental phenotype mostly similar to highly proliferative, disseminated tumor cells. Transitional gene expression signatures predict poorer patient prognosis in large cohorts of neuroblastoma. Moreover, cell type abundance of PNT microenvironments differ between neuroblastoma and other PNT subtypes. Finally, we use ligand-receptor modelling to infer cell-cell signaling relationships that indicate malignant cell subtypes have potential for distinct paracrine responses from their surrounding microenvironment.

## Results

### Single-cell transcriptomics analysis of peripheral neuroblastic tumors

We conducted single cell RNA sequencing using a modified 3’ unique molecular identifier (UMI) based version of the Smart-seq2 protocol^9^ on 7 peripheral neuroblastic tumors (Supplementary Fig. 1A). We used viable single cells (DRAQ5+/Calcein Blue+) derived from 3 neuroblastomas, 3 ganglioneuroblastomas and 1 ganglioneuroma (Supplementary Fig. 1B). Samples were acquired from surgical resection and had a range of clinical and histological features (Fig. 1A, Supplementary Table 1). Following sequencing, the median read depth was 22648 UMIs per cell with 1254 unique genes detected per cell (Supplementary Fig. 1C-E). We used quality control measures, such as the number of detected genes/cell (600-5000 genes per cell, where a given gene had to be expressed in at least 3 cells) and proportion of ribosomal counts (<0.3), to filter out poor quality cells (Supplementary Fig. 1C-F). Ultimately, we yielded 4267 high quality single-cell transcriptomes for downstream analysis across the 7 PNTs (range: 239-976 single cells per tumor). Using 4000 highly variable genes, we implemented principal component analysis (PCA) and selected the top 10 principal components for graph-based cell clustering (Supplementary Fig. 1G-H). This analysis identified 12 distinct clusters (Clusters 0 – 11) which was projected by uniform manifold approximation and projection (UMAP) (Fig. 1B-C).

**Figure 1:**
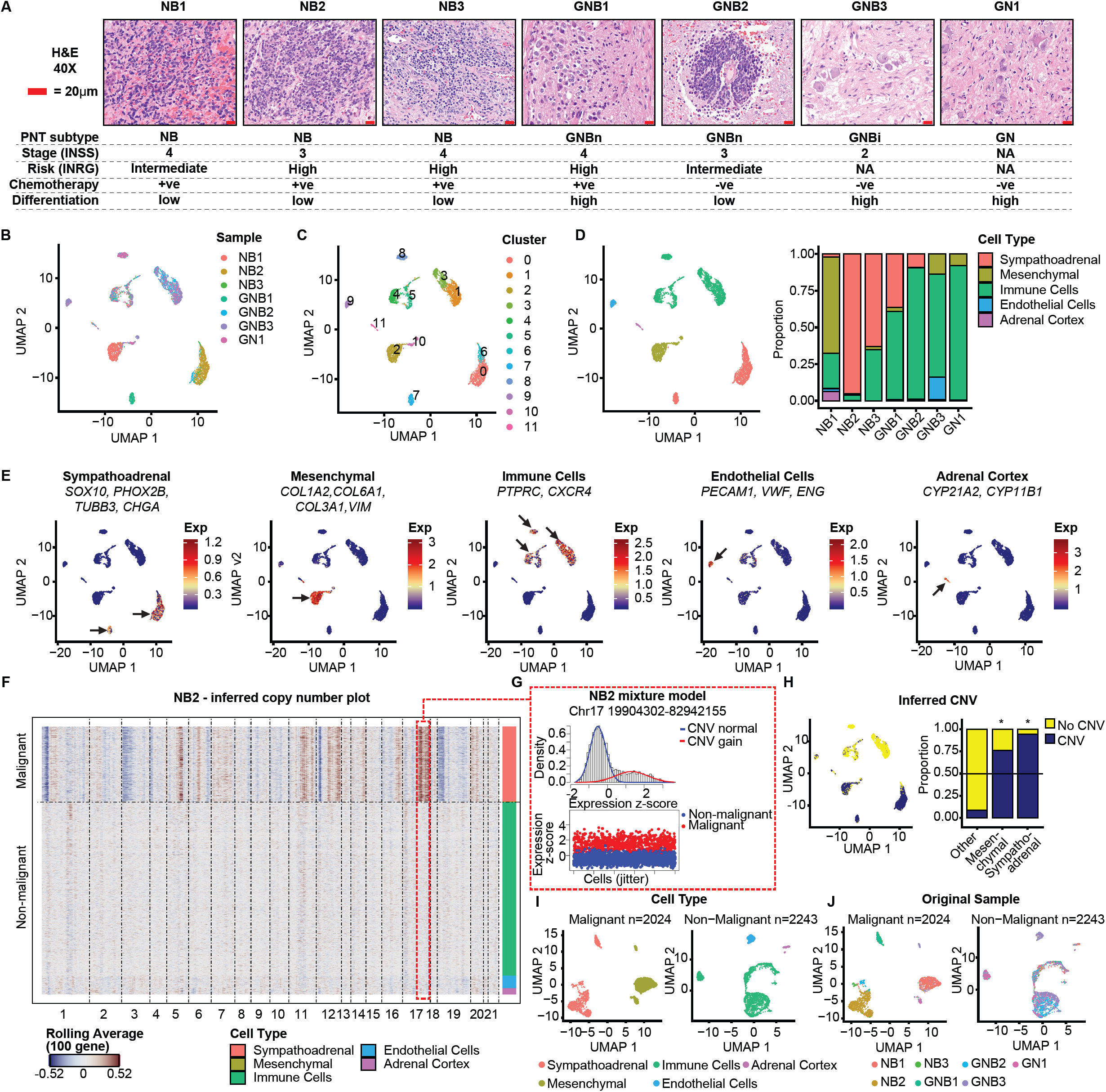
Single cell RNA-seq of peripheral neuroblastic tumors. A) Representative hematoxylin and eosin (H&E) staining in neuroblastoma (n=3), ganglioneuroblastoma (n=3) and ganglioneuroma (n=1) tumor samples. Scale bar: 20μm. Neuroblastomas are mostly composed of poorly/un-differentiated neuroblasts, which are small round cells with large nuclei. Ganglioneuroblastoma/ganglioneuroma are defined by the presence of mature ganglion cells and Schwannian stroma. B) Uniform Manifold Approximation and Projection (UMAP) plots of 4267 cells from all 7 tumor samples colored by original sample identity. C) UMAP plots of 4267 cells from all 7 tumor samples colored by transcriptomic cluster identity. D) Left: UMAP plots of 4267 cells classified by broad cell type based on marker gene expression from Fig. 1E. Right: Proportion of broad cell types within each of the 7 tumor samples. E) UMAP plots of 4267 cells from all 7 tumor samples. Average expression of marker genes for particular cell types (blue: low, dark-red: high). Marker genes and their associated cell types are indicated above each plot. Arrows indicate regions of high expression. F) Heatmap of CNVs for individual cells (rows), visualized by a rolling gene expression average centered on 100 gene windows at each chromosomal position (columns). Dark-red: copy number gains, blue: copy number losses. Colored sidebars denote cell type classification from Fig. 1D. Malignant cells (above dashed line) display an aberrant copy number profile compared to non-malignant cells (below dashed line). This plot uses the malignant cells from the NB2 sample compared with the non-malignant cells from all tumors. For a similar comparison for all other samples, see Supplementary Fig. 1L. G) Top: Two component Gaussian Mixture Models (GMM) were fit based on expression of genes located within the tested chromosome segments for each cell and tumor sample. Gaussian mixture components are show in different colors and overlaid on a density plot of 100-gene rolling average expression z-scores along the chromosome segment. A CNV was called in the component which had the highest proportion of inferred malignant cells. Significance of a segmental CNV call was determined for mixture models more likely to represent 2 components using Bayesian Information Criterion (BIC) metrics. See Supplementary Table 2 for all candidate CNV regions assessed this way and their respective BIC metrics. Bottom: Scatter plot visualization for each cells 100-gene rolling average in the tested CNV region. Malignant cells (red) and non-malignant cells (blue) are shown. Shown in this plot is a representative mixture model for Chromosome 17 in NB2, showing strong concordance between CNV values and predicted malignant cells. H) Left: UMAP plot of all cells (n=4267) colored by CNV classification. Confident CNV inferences were made for cells with a CNV score > 0.12. For CNV scores projected to a UMAP plot see Supplementary Fig. 1K. Right: Proportion of cells within each cell type based on CNV classification. If a cell cluster had >50% of cells classified to contain CNVs, then all cells in that cluster were classified as malignant (* indicates malignant cell types). I) UMAP plots showing preliminary cell type classification of: left: malignant cells (n=2024) or right: non-malignant cells (n=2243) split by CNV classification. J) UMAP plots showing original sample identity of: left: malignant cells (n=2024) or right: non-malignant cells (n=2243) split by CNV classification.

To assign each cluster to a cell phenotype, we undertook a preliminary sub-classification that considers differential expression and the expression of marker genes for major cell lineages (Fig. 1D-E, Supplementary Fig. 1I-J). Based on these cell-type assignments, the proportion of cell types within individual tumors differed markedly (Fig. 1D). Notably, most clusters conformed to broad lineage classifications that would be expected to occur in PNTs^7,8,10-15^, such as sympathoadrenal, immune, mesenchymal, endothelial and adrenal cell types (Fig. 1D-E).

### Identification of malignant neuroblasts by copy number variation inference

With the knowledge that malignant cells in neuroblastic tumors can exist in multiple differentiated forms^7,8^, classification of malignant cells using gene expression alone may be inaccurate. We therefore aimed to distinguish malignant cells from non-malignant cells using CONICS copy number estimation as an additional method of classification^16^. This technique uses the expression of a sliding window of 100 genes across each chromosome to infer copy number values (CNVs), so that malignant cells with greater copy number instability can be identified (Fig. 1F, Supplementary Fig. 1L). Two-component Gaussian mixture models were calculated to identify single cell copy number estimations for various regions of the genome, using the cells of each tumor (for example Fig. 1G, Supplementary Table 2). From these models, a CNV score was calculated to approximate whether a cell had CNV’s detected for these individual regions (see methods, Supplementary Fig. 1K). Projection of inferred CNV’s on the transcriptomic UMAP plot showed that the CNV distribution was non-random and localized to specific clusters of cells (Fig. 1H, Supplementary Fig. 1K). Indeed, the proportion of cells inferred to contain a CNV showed that sympathoadrenal and mesenchymal clusters consisted mainly of the cells with CNVs, while other clusters had relatively few CNVs (Fig. 1H). Heatmap visualization of 100 gene rolling averages for each cell similarly showed greater copy number instability in these specific cell types (Fig. 1F, Supplementary Fig. 1L). Using these inferred CNV’s, we therefore concluded that clusters subclassified as sympathoadrenal or mesenchymal were malignant cells while all other clusters were non-malignant cells (Fig. 1I). After splitting the cells into malignant (n=2024) and non-malignant (n=2243) groups, UMAP projection of malignant cells separated mostly into distinct subpopulations associated with individual patient tumors, suggesting pronounced intertumoral heterogeneity among neuroblasts (Fig. 1J, left panel). In contrast, non-malignant cells tended to cluster independent of tumor origin (Fig. 1J, right panel), consistent with previous reports showing non-malignant cells cluster by cell type rather than the tumor that they derive from^17,18^.

### Modelling of adrenergic/mesenchymal transdifferentiation identifies a novel transitional cell phenotype

To evaluate cell phenotype among malignant cells, we conducted differential expression analysis and identified divergent cell transcriptomes highlighted by expression of adrenergic (CHGA, CHGB, DBH) and mesenchymal marker genes (COL1A1, COL8A1, NOTCH3) (Supplementary Fig. 2A-B). We therefore investigated the possibility that malignant cells exist in either an adrenergic/noradrenergic or mesenchymal/neural crest cell state, as proposed by recent reports by van Groningen and Boeva *et al*^*7,8*^. We first examined expression patterns of signatures for these cell states. UMAP projection of these two gene signatures showed expression that was enriched in distinct clusters whether using the van Groningen (Adrenergic/Mesenchymal) or Boeva signatures (noradrenergic/neural crest cell) (Fig. 2A-B, Supplementary Fig. 2C-D). When comparing directly, the expression patterns of adrenergic and mesenchymal signatures showed a strong inverse association, with cells existing in a continuum between high-expressing adrenergic and mesenchymal cells (Fig. 2C). This suggests that cells may have an identity that reflects various points of transdifferentiation between adrenergic and mesenchymal cell states, similar to the model proposed by van Groningen and Boeva *et al*^*7,8*^. To explore this hypothesis further, we conducted pseudotemporal ordering of cells^19^ to model cell transitions that would occur during adrenergic-mesenchymal transdifferentiation (Fig. 2D). Interestingly, the predicted trajectory was more complex than a simple linear path running between adrenergic and mesenchymal cell states. Examination of the adrenergic and mesenchymal signatures showed a unique state as a separate arm between the high adrenergic and high mesenchymal expression states, which we refer to as “transitional” cells (Fig. 2E-G). Gene expression analysis of this novel transitional cell state demonstrated an intermediate expression level between adrenergic and mesenchymal signatures that mostly corresponded to cluster 4 cells but also some cluster 2 cells (Fig. 2H). We also considered whether cells derived from each tumor contained cells from more than one neuroblastic state, as would be expected if cells had the capacity to transdifferentiate. Indeed, most tumors had evidence of cells in more than one neuroblastic state, except the lower risk GNB3 and GN1 tumors (Fig. 2I, Supplementary Fig. 2E-F). These data suggest that extensive cell phenotype heterogeneity exists in PNTs, with cell state corresponding to various points along a adrenergic-mesenchymal transdifferentiation trajectory.

**Figure 2:**
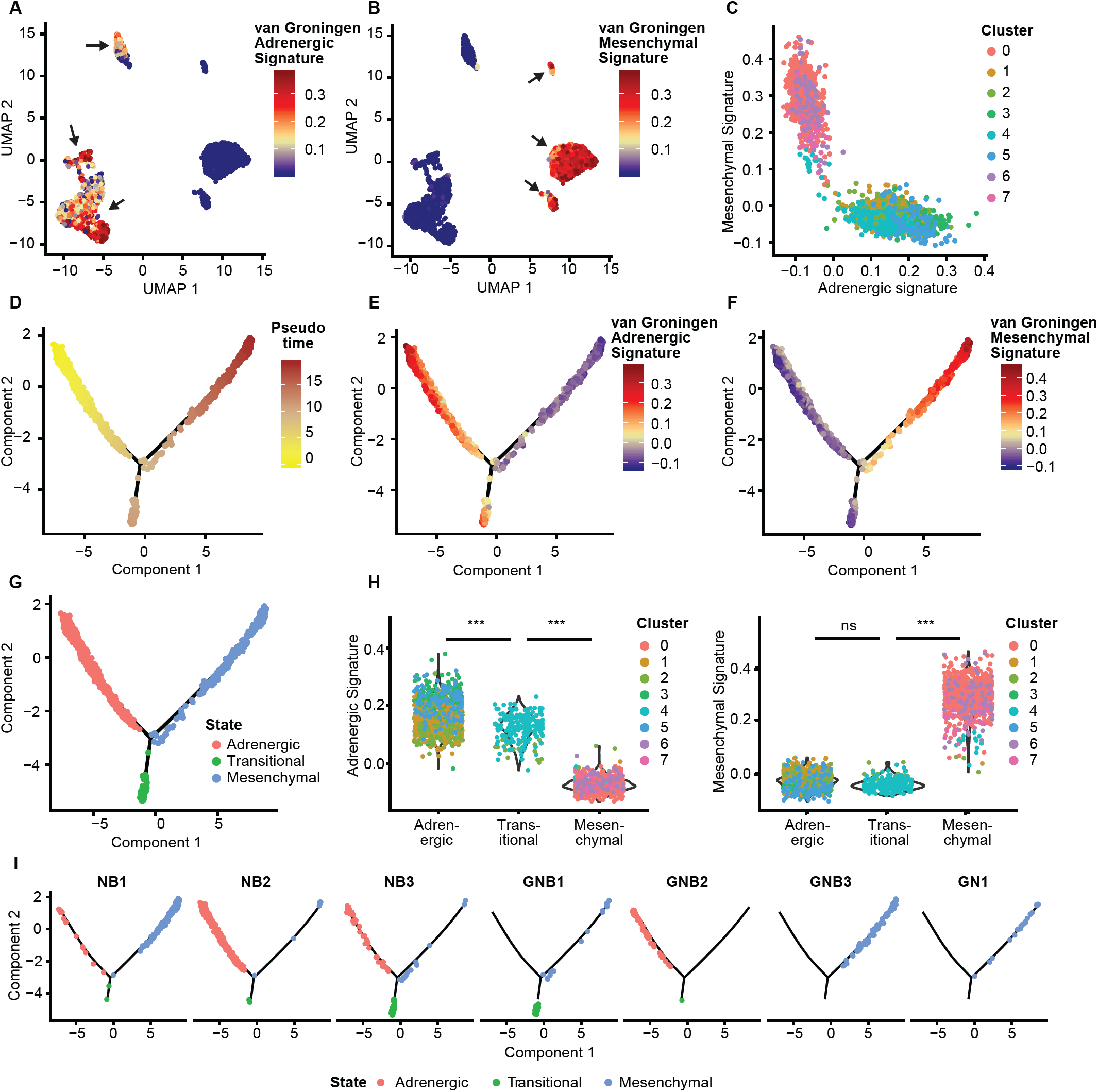
Trajectory modelling of malignant neuroblasts identifies a transitional phenotype as an intermediate state between adrenergic and mesenchymal states. A) UMAP of 2024 malignant cells colored based on the average expression of genes in a previously published adrenergic gene signature^8^ (blue: low, dark-red: high). B) UMAP of 2024 malignant cells colored based on the average expression of genes in a previously published mesenchymal gene signature^8^ (blue: low, dark-red: high). C) Scatter plot of malignant cells (n=2024) based on adrenergic and mesenchymal gene signature expression. Cells colored according to transcriptomic cluster defined in Supplementary Fig. 2A. D-G) Trajectory-based inference of malignant cells (n=2024) in a two-dimensional space using Monocle2 (DDRTree). Cells are represented as individual dots ordered along the trajectory (solid lines) by increasing pseudotime^19^. Pseudotime was modelled based on expressed genes from previously defined signatures^8^ (see methods). (D) Cells are colored according to pseudotime using the left most arm as the root of the trajectory. (yellow: low, brown: high). (E) Cells are colored according to the average expression of genes within the adrenergic gene signature^8^ (blue: low, dark-red: high). F) Color indicates the average expression level of genes within the mesenchymal gene signature^8^ (blue: low, dark-red: high). (G) Cells colored by inferred cell state (adrenergic (n=1007), transitional (n=209) or mesenchymal (n=808)). H) Violin plots of adrenergic (left) and mesenchymal (right) gene signatures for all malignant cells segregated by inferred cell state in Fig. 2G. Cells are colored according to transcriptomic cluster defined in Supplementary Fig. 2A. P-values are provided for comparisons made using t-test (***, p<0.001). I) Trajectory plots for each tumor sample, illustrating which cell state (in Fig. 2G) cells belong to. Cells are colored by cell state identity.

### Transitional neuroblasts are defined by an aggressive neurodevelopmental phenotype

In order to clarify transcriptomic change during cell state transition, we identified marker genes in each of the 3 cell states, adrenergic, transitional and mesenchymal (Supplementary Fig. 3A-B, Supplementary Table 3). Evaluation of changes in gene expression using pseudotime to visualize cell trajectories, showed that the expression of these genes progressively changed when comparing different states (Fig. 3A, Supplementary Fig. 3C). Key noradrenergic and catecholaminergic enzymes expressed in the adrenergic state, dopamine beta-hydroxylase (DBH) and tyrosine hydroxylase (TH), were down-regulated along trajectories to either the transitional or mesenchymal arm (Fig. 3A, top panel). The mesenchymal state showed upregulation of mesenchymal neuroblast markers FN1 and VIM, consistent with a role in extracellular matrix production expected in this cell type (Fig. 3A, bottom panel). In transitional neuroblasts, we found an overexpression of well-known neuroblastoma genes EZH2 and MYCN (Fig. 3A, middle panel). While MYCN is an established driver gene in the context of neuroblastoma when gene-amplified^20^, this was an interesting finding since MYCN is non-amplified in all the tumors of this cohort. EZH2 on the other hand is a core component of polycomb repressive complex 2 and responsible for the catalysis of H3K27 tri-methylation. EZH2 plays an essential role in tumorigenesis in neuroblastoma and aberrant expression of EZH2 is strongly associated with poor prognosis^21-23^. Next, we employed gene ontology (GO) analysis on gene signatures created for each of the three cell states. GO-term enrichment identified gene-sets as expected related to neurotransmitter production and extracellular matrix as part of the adrenergic and mesenchymal states, respectively (Fig. 3B, Supplementary Fig. 3D-E). Transitional neuroblasts, however, had gene-sets relating to neurogenesis (Fig. 3B, Supplementary Fig. 3D-E), suggesting that differentiation via transitional neuroblasts may be a developmentally co-opted process. Indeed examination of single cell RNA-seq data in E12.5 murine neural crest compartment^24^, shows that the transitional signature shows relatively high expression in the so-called “bridge” cells that act as differentiation intermediates between the Schwann cell precursors of the neural crest (mesenchymal-like) and the terminal chromaffin cells of the adrenal medulla (adrenergic-like) (Fig. 3C, Supplementary Fig. 3F).

**Figure 3:**
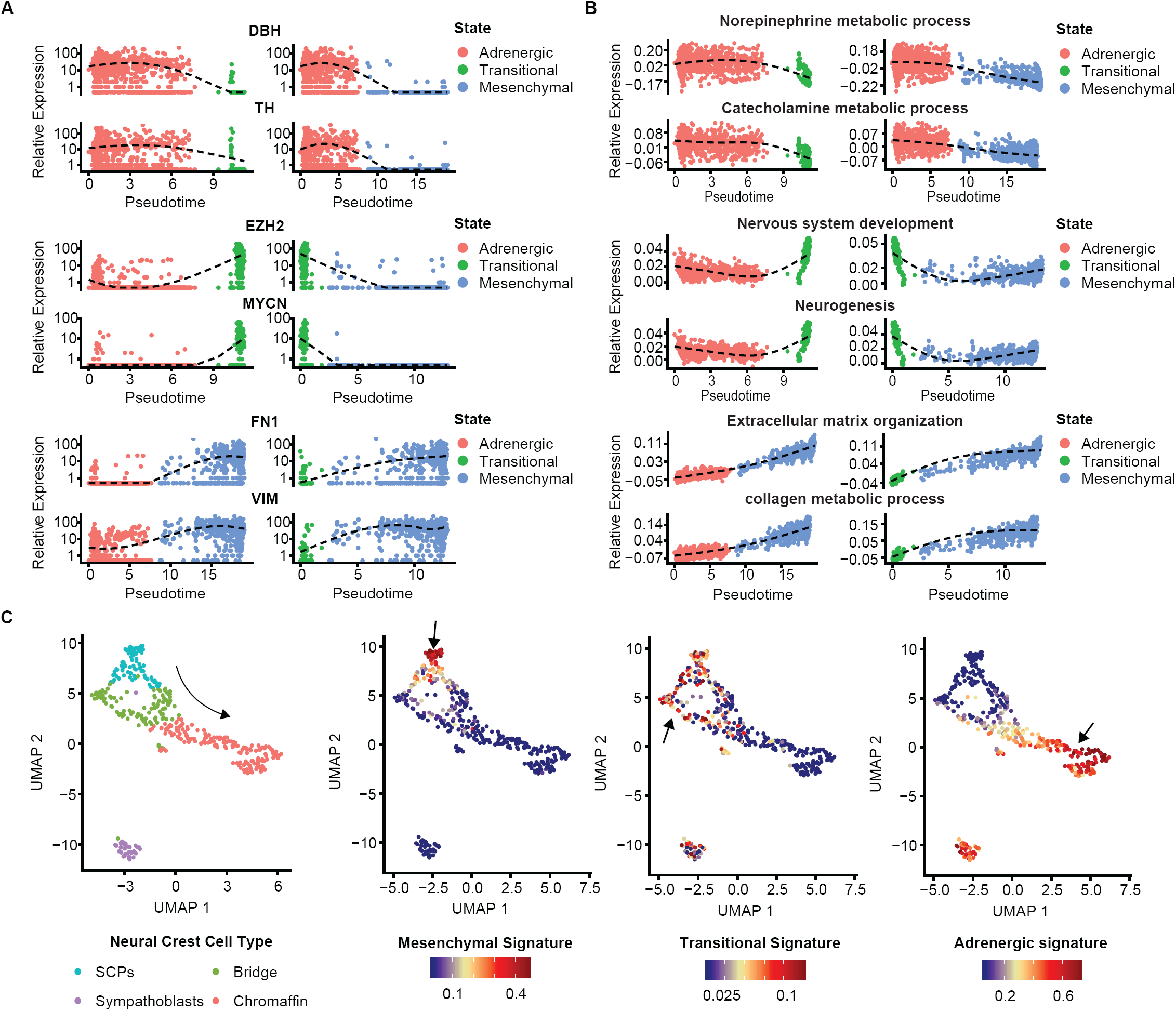
Malignant neuroblasts in PNTs have gene expression characteristics representative of various differentiated stages in the developing neural crest. A) Gene expression of individual cells ordered by pseudotime along arms of the trajectory in Fig. 2G. Each point represents a cell, with the dashed line representing a natural cubic splines curve fitted to all points. Pseudotime was calculated considering the adrenergic to transitional trajectory, the adrenergic to mesenchymal trajectory or the transitional to mesenchymal trajectory depending on the respective “states” featured in each plot. Marker genes which were significantly altered along both trajectory arms are used as examples; adrenergic (DBH, TH), transitional (EZH2, MYCN) and mesenchymal (FN1, VIM). Cells are colored according to cell states. B) GO signature expression of individual cells ordered by pseudotime along arms of the trajectory in Fig. 2G. Each point represents a cell, with the dashed line representing a natural cubic splines curve fitted to all points. Pseudotime was calculated considering the adrenergic to transitional trajectory, the adrenergic to mesenchymal trajectory or the transitional to mesenchymal trajectory depending on the respective “states” featured in each plot. Marker pathways which were significantly altered along both trajectory arms are used as examples; adrenergic (norepinephrine metabolic process, catecholamine metabolic process), transitional (nervous system development, neurogenesis) and mesenchymal (extracellular matrix organization, collagen metabolic process). Cells are colored according to cell states. C) UMAP plots on murine neural-crest cells from embryonic day 12.5. Left: Shows the major cell types as defined in Furlan *et al*^24^. Right: Expression of adrenergic, transitional and mesenchymal state signatures in neural crest cells.

We next investigated whether the three cell states related to biological processes important in tumorigenesis. To gain more insight into the cell cycle status of the malignant cells across the different phenotypes, we determined their cell cycling states based on the average expression levels of genes within the S and G2/M (dividing) and G1 (non-dividing) gene sets previously published^18^. This analysis revealed a markedly higher proportion of dividing cells amongst the transitional neuroblasts compared to the other two cell states (Supplementary Fig. 3G). In contrast, mesenchymal and adrenergic cells had a greater proportion of non-dividing cells with expression of the G1 phase-related genes (Supplementary Fig. 3G). Next, we evaluated whether either of the three cell states related to a metastatic phenotype. For this, we calculated a disseminated tumor cell (DTC) score for gene expression in each cell state by determining the difference between the expression of significantly up-regulated and down-regulated genes of a previously published DTC gene set^25^. Interestingly, the transitional malignant cells shared the highest similarity with DTCs from the bone marrow which is the most common distant metastatic site of neuroblastoma (Supplementary Fig. 3H). Consistent with this, tumors with a high abundance of transitional neuroblasts, NB3 and GNB1, were graded as Stage 4 with distant metastasis by the International Neuroblastoma Staging System (INSS), further suggesting a link between the transitional phenotype and metastasis. Collectively, these data suggest the presence of distinct malignant cell subpopulations within individual tumors, which demonstrate divergent differentiation status, varying transcriptional signatures and potential for malignant clinical behavior.

### Tumors expressing transitional neuroblast signatures are associated with poor prognosis in neuroblastoma patients

We next studied expression data from large primary neuroblastoma tumor cohorts where gene expression was derived from total tumor or bulk gene expression data (Kocak cohort, n=649; GEO GSE45480)^26^.We classified tumors based on adrenergic, transitional and mesenchymal gene signatures (Fig. 4A). The signatures separated 5 tumor groups, ranging from adrenergic, transitional and mesenchymal, as well as two mixed classes of adrenergic-transitional, and transitional-mesenchymal states (Fig. 4A). This is consistent with the notion that in some cases, bulk tumors can have mixed proportions of the three cells states identified in our scRNA-seq data. To explore whether the three cell states had prognostic relevance, we conducted Kaplan-Meier analysis for the 5-tumor subgroups defined by these signatures. Transitional tumor classes were shown to predict poorer outcome than adrenergic and mesenchymal subgroups, particularly the transitional-only and transitional-mesenchymal subgroup (Fig. 4B). Additionally, higher expression of transitional neuroblast signatures was observed in patient subgroups defined by poor prognostic factors: *MYCN-*amplification and INSS 4 disease (Fig. 4C). This suggests that transitional cell state signatures identify tumors with high risk of resistance to conventional therapy and relapse. Consistent with these findings, Kaplan-Meier analyses for gene signatures subdividing the cohort at the median showed a similar association between transitional neuroblasts and poor prognosis, as did similar analyses in a separate cohort (Versteeg cohort, n=88; GEO GSE16476)^27^ (Supplementary Fig. 4A-E).

**Figure 4:**
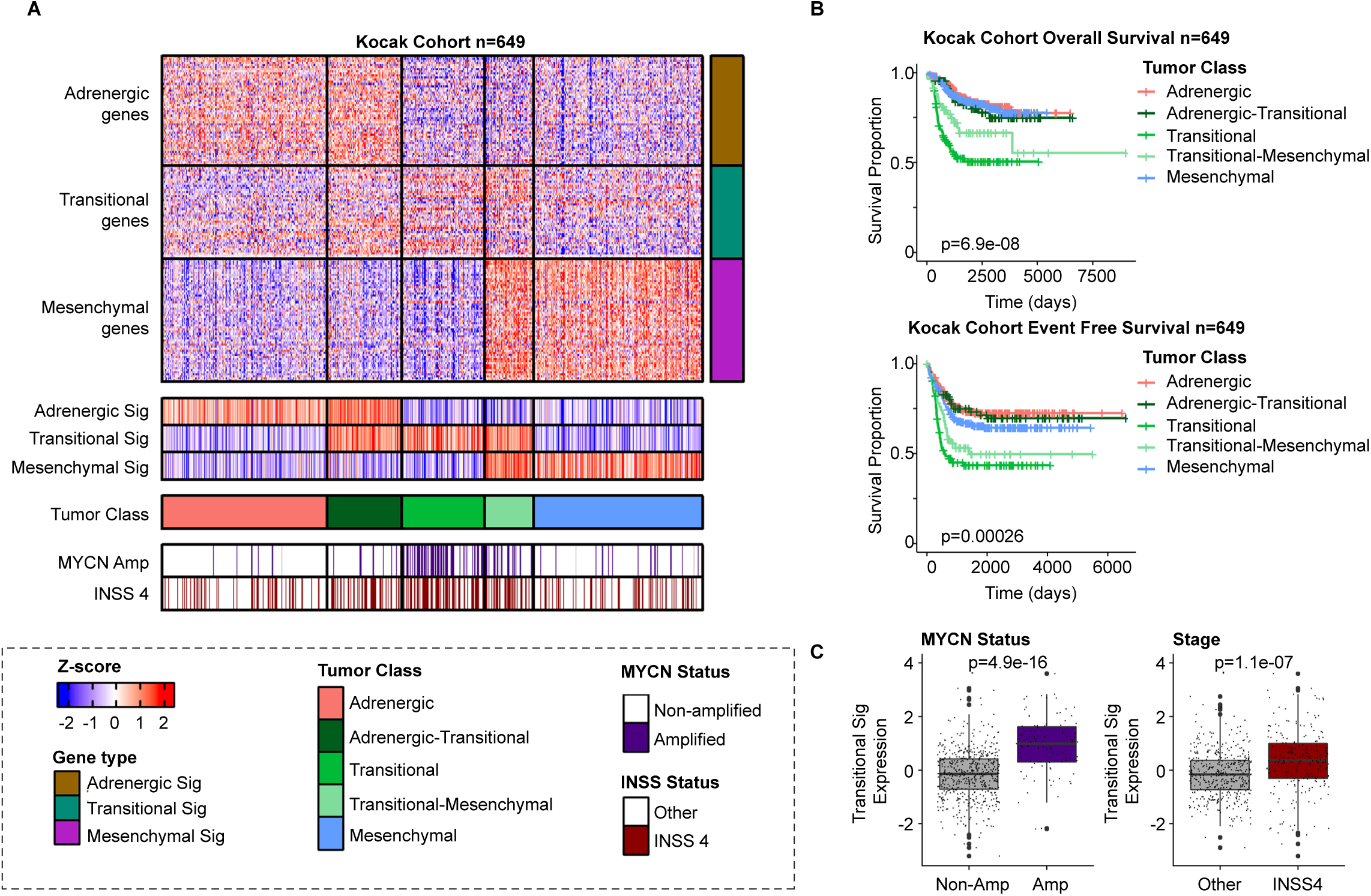
Tumors expressing transitional neuroblast signatures are associated with poor prognosis in neuroblastoma. A) Heatmap detailing the classification of neuroblastoma patients from a bulk mRNA (microarray) profiling cohort (Kocak, n=649 patients^26^) based on adrenergic, transitional and mesenchymal gene signatures. Gene expression values (rows) for each tumor (columns) were arranged based on their respective gene signature. Column averages were used for each signature (shown immediately below the heatmap) to classify tumors to Tumor Class subgroups: Adrenergic-only, Adrenergic-Transitional, Transitional-only, Transitional-Mesenchymal, and Mesenchymal-only (see methods). Patient metadata concerning; *MYCN* status (amplified) and neuroblastoma staging (INSS 4) are annotated in bars underneath the heatmap. Scale bar indicates gene expression z-scores. B) Kaplan-Meier plots of overall survival (top) and event free survival (bottom) in the Kocak cohort colored based on Tumor Class classifications made in Fig. 4A. P-value was calculated using log-rank tests, comparing all transitional subgroups combined to adrenergic and mesenchymal combined. C) Boxplots of transitional gene signature expression in patients of the Kocak cohort (n=649), when dichotomized by MYCN amplification status (left; non-amplified vs amplified) or INSS staging (right; stage 1, 2, 3 & 4S vs stage 4). P-values are reported from 2-sided t-tests, and boxes represent first-quartile/median/third-quartile of expression values, with the whiskers representing the 95% confidence interval, and large points representing outliers. Small points represent individual tumors.

### Neuroblastoma cell lines with high transitional gene signature expression have an intermediate super-enhancer landscape between adrenergic and mesenchymal states

Previous studies used H3K27-acetylation (H3K27ac) landscapes to define two distinct super-enhancer states, either adrenergic/noradrenergic or mesenchymal/neural crest^7,8^. Since our trajectory analysis using gene expression in single-cells identified a transitional population between these two identities, we investigated whether transitional gene expression signatures in neuroblastoma cell lines, similarly identify intermediate super-enhancer profiles. As before, we used adrenergic, transitional and mesenchymal signatures to classify 33 neuroblastoma cell lines with paired bulk-RNA-seq and H3K27ac ChIP-seq data^7^ into 5 groups: either adrenergic, adrenergic-transitional, transitional, transitional-mesenchymal or mesenchymal (Supplementary Fig. 5A). We then conducted comparisons to identify super-enhancers that were enriched in either the adrenergic class or the mesenchymal class (Supplementary Table 4A). Of the 5975 previously annotated super-enhancer loci across these 33 cell lines^7^, we found 1129 were enriched in the adrenergic state (versus mesenchymal cell lines), and 2203 were enriched in the mesenchymal state (versus adrenergic cell lines) (Fig. 5A, Supplementary Fig. 5B). To evaluate if a progressive shift in H3K27ac occurs through cell lines according to our cell state classification, we investigated if adrenergic or mesenchymal-associated superenhancers showed any trend across different cell classes. H3K27ac levels at adrenergic-associated super-enhancers progressively decreased through the spectrum of adrenergic, adrenergic-transitional, transitional, transitional-mesenchymal and mesenchymal cell lines (Fig. 5A - left panel, Fig. 5B, Supplementary Fig. 5B - upper panel, Supplementary Fig. 5C). In contrast, H3K27ac levels at mesenchymal-enriched super-enhancers progressively increased through the same spectrum (Fig. 5A – right panel, Fig. 5B, Supplementary Fig. 5B - lower panel, Supplementary Fig. 5C). Moreover, H3K27ac tracks show a similar trend in super-enhancers linked to core-regulatory circuit transcription that were previously found to define adrenergic and mesenchymal identities^7^ (Fig. 5C). Interestingly, transitional-enriched super-enhancers did exist, albeit only in the minority, with only 94 unique super-enhancers significantly enriched in transitional cells compared with adrenergic and mesenchymal cells (Supplementary Fig. 5D, Supplementary Table 4B). There were however some examples of super-enhancer loci significantly upregulated exclusively in transitional cell lines (Supplementary Fig. 5E, Supplementary Table 4B). These results suggest that indeed, expression of transitional cell state signatures in neuroblastoma cell lines was associated with intermediate super-enhancer profiles that lie between adrenergic/nor-adrenergic and mesenchymal/neural crest cell differentiation states. Notably however, these results show that while transitional cells lie between adrenergic and mesenchymal cell states, they tended to have a H3K27ac profile toward the adrenergic end of the transdifferentiation spectrum (Fig. 5A-B).

**Figure 5:**
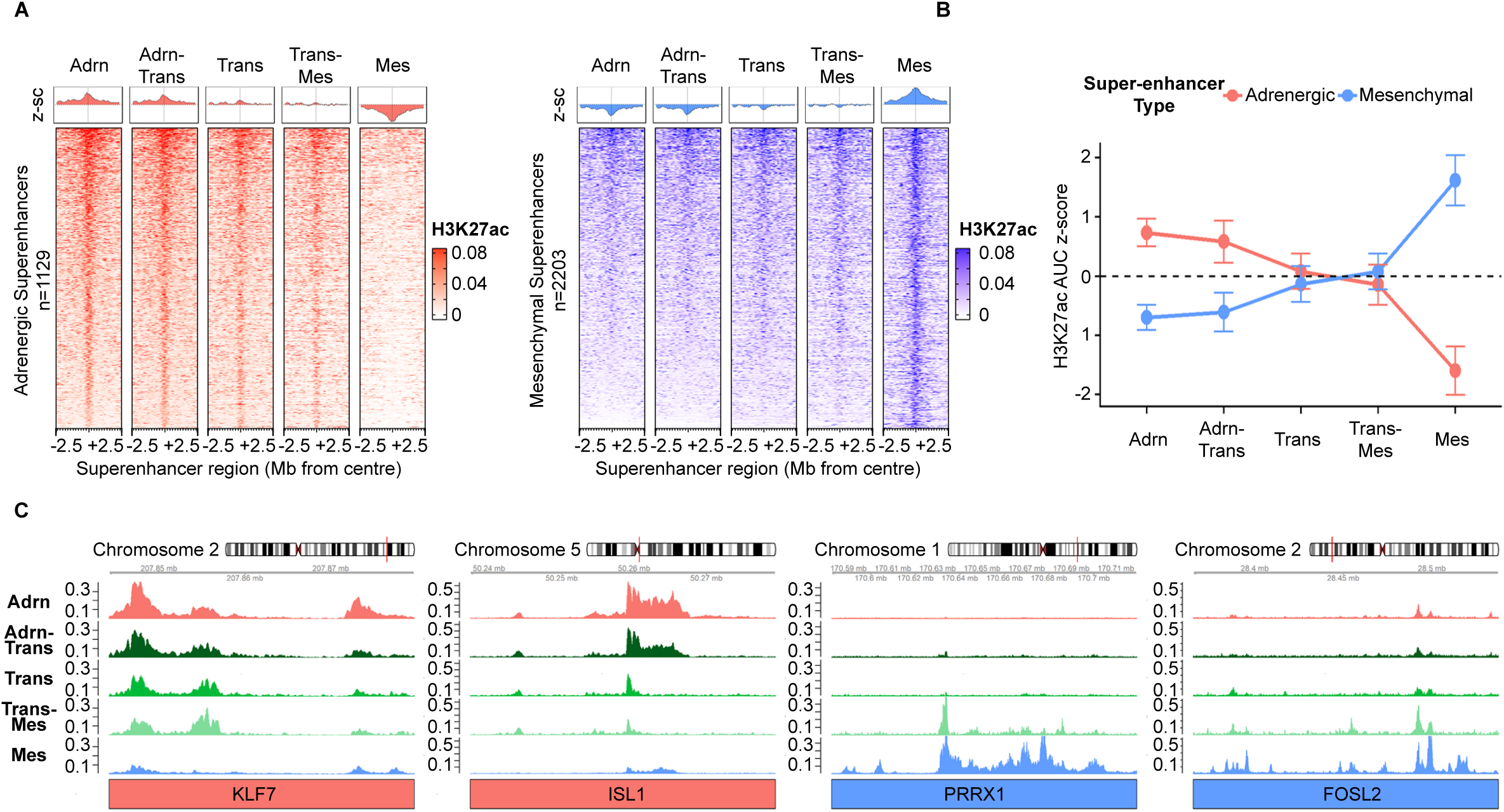
Neuroblastoma cell lines expressing transitional neuroblast signatures are at the junction between adrenergic and mesenchymal super-enhancer states. A) Global normalized H3K27-acetylation (H3K27ac) values for cell lines classified to 5 subgroups of cells lines as listed (see, Supplementary Fig. 5A for gene expression classification). Super-enhancer loci enriched in either the Adrenergic cell lines (left panel, red), or Mesenchymal cell lines (right panel, blue) are shown. H3K27ac values are averaged across all cell lines classified in that group and represent 0.5Mb centered on the super-enhancer locus in that region. For individual cell line traces, see Supplementary Fig. 5B. Above each set of traces is the average H3K27ac z-score value for that binned region of the trace for each respective type of enriched super-enhancer (Adrenergic super-enhancer H3K27ac average z-score: red, Mesenchymal super-enhancer H3K27ac average z-score: blue). Bin size for each trace is 1kb. B) Global normalized H3K27ac values for adrenergic superenhancers (red) and mesenchymal super enhancers (blue) were quantified for each of the classified 5 subgroups of cell lines using a z-score value of the total area under the H3K27ac curve. Error bars represent the standard deviation of H3K27ac AUC z-score for the individual cell lines allocated to each of the 5 tumor class subgroups. C) Representative H3K27ac for each group of classified cell lines for core-regulatory circuit transcription factor (CRC TFs) super-enhancer loci. Red are examples of adrenergic CRC TFs and blue are examples of mesenchymal CRC TFs.

### Neuroblastomas and ganglioneuroblastomas have distinct tumor microenvironmental cell types

Next, we examined gene expression characteristics of cells in the PNT tumor microenvironment. The proportion of infiltrating immune cells was higher in the GNB and GN tumors when compared to the three neuroblastoma samples (Fig. 6A). When we excluded malignant cells and interrogated cell-type abundance in each tumor as a proportion of all non-malignant cells we found nine distinct cell clusters (Fig. 6B). Based on differential expression analysis and known markers of cell type, we then classified cells into subtypes, and found that main clusters conformed to recognized cell types from the immune, adrenal, or endothelial lineages (Fig. 6C-D, Supplementary Fig. 6A-C). When comparing the relative abundance of different cell types, the three neuroblastomas had less total infiltrating T-cells, including cytotoxic T-cells (measured by CD8 and presence of cytolytic effector genes) than the non-neuroblastoma samples (Fig. 6D, Supplementary Fig. 6B-C – upper panels). Consistent with these findings, immunohistochemistry of tissue sections from each of the seven tumors showed that pan-T (CD3) positive cells were more abundant in GNB2, GNB3 and GN samples compared with neuroblastoma (Supplementary Fig. 6D). Evaluation of other cell types revealed that neuroblastoma samples had more macrophages, including non-inflammatory macrophages with increased expression of the M2-polarization marker CD163 (Fig. 6D, Supplementary Fig. 6B-C – lower panels). Previous studies have revealed that tumor-associated macrophages can be induced by neuroblastoma cells and polarized into an M2/pro-tumor phenotype, which then can promote the proliferation and invasion of tumor cells^28,29^. Our analysis of the cellular makeup of the tumor microenvironment showed a trend of increased T-cell infiltration in ganglioneuroblastomas and increased macrophage infiltration in neuroblastoma. This suggests a potential role of immune evasion and a pro-tumorigenic microenvironment in the more aggressive neuroblastomas, comparing with non-neuroblastomas.

**Figure 6:**
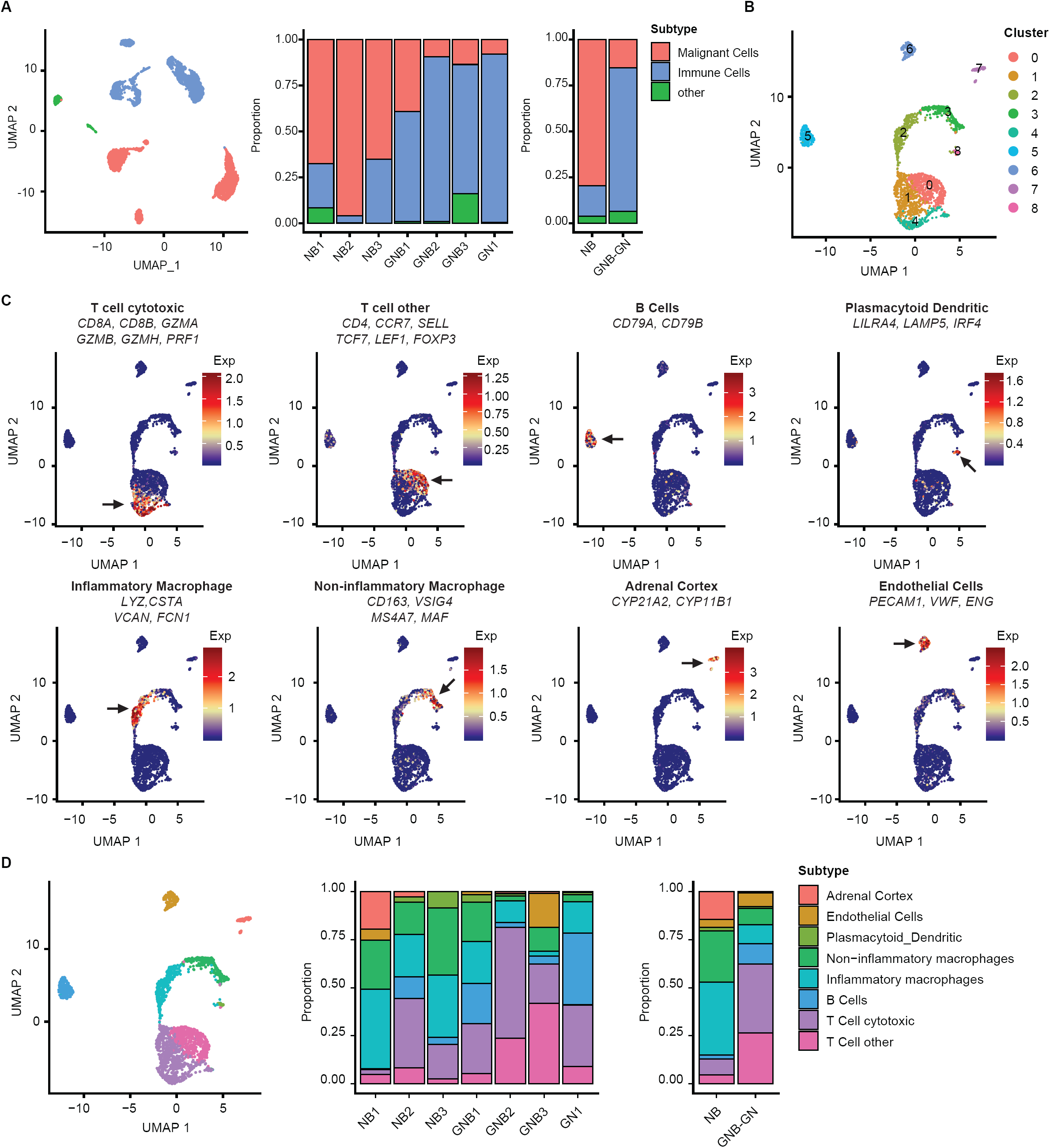
Neuroblastomas and ganglioneuroblastomas have distinct tumor microenvironmental cell types. A) UMAP plot showing malignant (n=2024), immune (n=2022) and other (n=221) cell types from 7 tumors. The proportion of cell types within each tumor and major tumor types are also shown (neuroblastoma compared with ganglioneuroblastoma/ganglioneuroma). Colors indicate the various cell types. B) UMAP plot of 2243 non-malignant cells from 7 tumors with cells colored based transcriptomic cluster. C) UMAP plot of 2243 non-malignant cells from 7 tumors with cells colored based on the average expression of marker genes for specific cell types (marker genes and associated cell types are indicated above to each plot). Scale: gene signature expression (blue: low, dark-red: high). Arrows indicate regions of high expression. D) Left: UMAP plot showing subtypes of non-malignant cells (n=2243) from 7 tumors. Right: the proportion of non-malignant cell types within each tumor and major tumor types are also shown (neuroblastoma compared with ganglioneuroblastoma/ganglioneuroma). Colors indicate the various non-malignant cell subtypes.

### Discovery of unique cell-cell signaling potential within tumor microenvironments

Upon resolving the complex tumor microenvironments of our PNT cohort, we considered whether we could use our data to infer cell-cell signaling mediated by non-malignant cells driving gene expression states in malignant cells. Based on the target gene signatures that define the adrenergic, transitional and mesenchymal states, we employed probabilistic models to infer cell-cell signaling networks in our scRNA-seq data using NicheNet^30^ (Supplementary Fig. 7A-D). NicheNet integrates known ligand-receptor and ligand-target interactions to infer signaling relationships based on correlation of gene expression. We modelled ligand-receptor-target interactions between non-malignant and malignant cell populations for each patient yielding 87 interactions, in which a ligand expressed by a specific non-malignant cell-type was predicted to bind a receptor and effect target gene expression in malignant cells (Fig. 7A, Supplementary Fig. 7E, Supplementary Table 5). When evaluating the amount of interactions predicted for each of the three malignant cell states we previously defined (i.e. adrenergic, transitional and mesenchymal), we found that the majority of predicted interactions occurred between non-malignant cells and mesenchymal neuroblasts (Fig. 7B). Non-malignant endothelial cells facilitated the majority of interactions with mesenchymal neuroblasts (Fig. 7B, Supplementary Fig. 7E-F), which is consistent with the migratory phenotype of mesenchymal neuroblasts and their reliance on vascular interactions preceding systemic dissemination^8,31^. Moreover, other specific cell-cell interactions were also found among the 3 malignant cell types. Interestingly, we found the expression of tumor associated transcriptional regulators such as ZFHX3, MYCN and NUPR1 to be correlated with specific ligand-receptor interactions in their respective malignant cell types, suggesting a role for cell-cell interactions in promoting broader transcriptional activity in subsets of malignant cells (Fig. 7C). Taken together, these data implies a role for the PNT tumor microenvironment in influencing different malignant cell states through cell-cell signaling, and highlights potential paracrine cell signaling pathways involved in PNT microenvironments.

**Figure 7:**
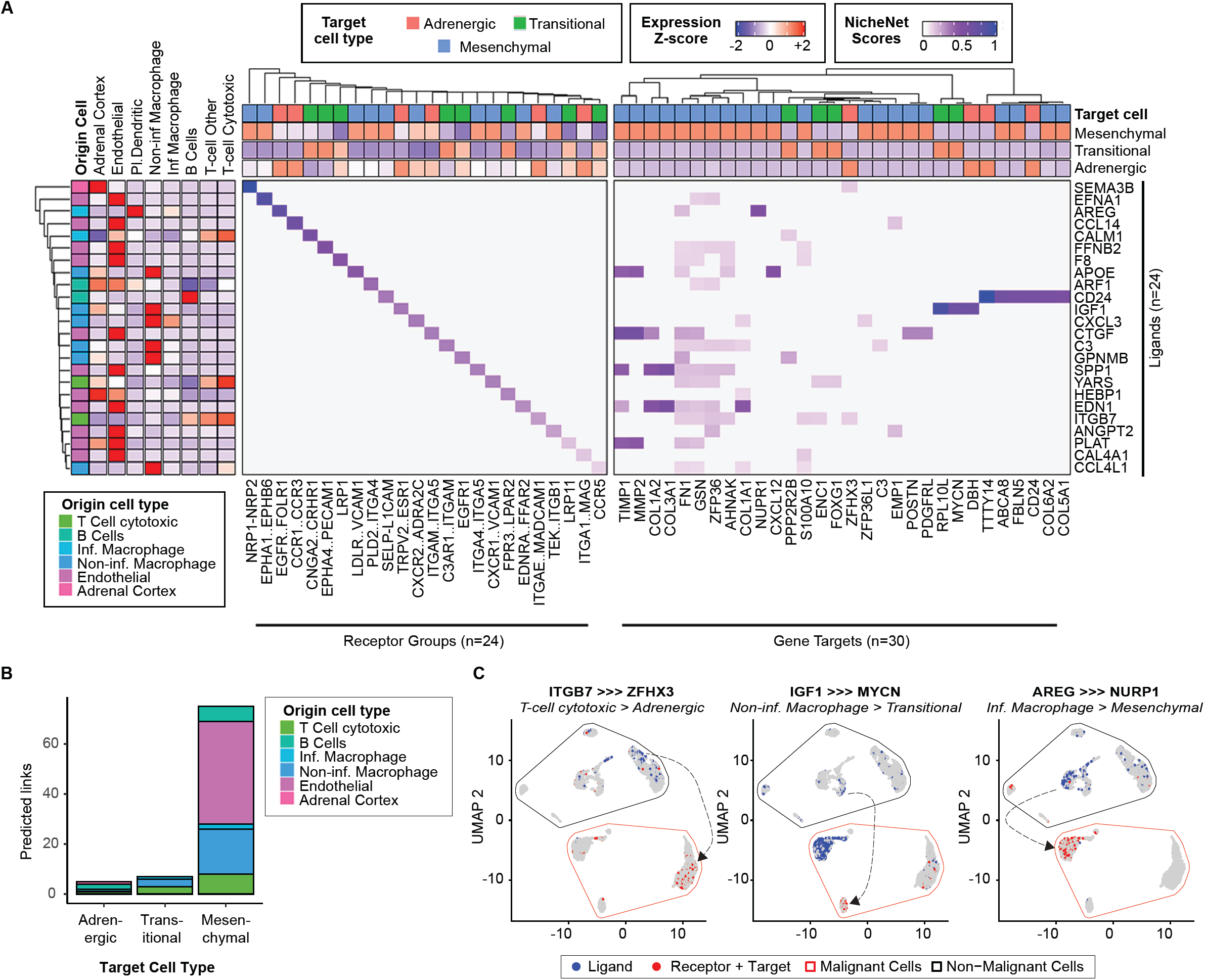
Modelling of cell-cell signaling in peripheral neuroblastic tumors. A) Heatmap representation of normalized ligand-receptor and ligand-target interaction scores from NicheNet models. Rows represent ligands from non-malignant cells (n=24) with predicted receptor groups (n=24) or gene targets (n=30) in malignant cells as columns. Cell-type specificity of the indicated ligands from predictions are given alongside scaled expression values of the ligand in non-malignant cell types. Specificity of gene targets/receptors are based on gene expression and are annotated alongside scaled expression values of each receptor group/target in malignant cell subsets. Scales of the interaction scores for ligand-receptor and ligand-target as well as scaled gene expression are provided above the heatmap. Rows and columns were hierarchically clustered using Euclidean distance and ward.D2 functions, clustering dendrograms are shown. B) Stacked column graphs showing the number of ligand-receptor-target links (n=87) between adrenergic, transitional and mesenchymal neuroblasts and non-malignant cells. Bars are colored by non-malignant cell type specificity; annotations are displayed to the right. C) UMAP plots of malignant and non-malignant cells (n=4267 cells), colored based on whether the cell expresses a given ligand (blue) or its respective receptor & target (red, indicates receptor and downstream target are expressed in same cell). Malignant & Non-Malignant cell groups are enclosed on the UMAP.

## Discussion

Patients with peripheral neuroblastic tumors (PNTs) are characterized by extensive inter-and intratumoral heterogeneity^5,32,33^. Recent studies have reported that PNTs were comprised of different types of transdifferentiating malignant cells with dissimilative epigenetic landscapes and gene expression profiles^7,8^. Moreover, the tumor microenvironment (TME) plays an important role in tumor biology and adds more complexity to intratumoral heterogeneity^34,35^. Here we provide comprehensive analysis of the cellular heterogeneity of PNTs through single-cell transcriptomic analysis of 4267 cells obtained from seven surgically resected tumors. Our data revealed 11 unique cell types, corresponding to 3 malignant cell types, 6 immune cell types, and 2 other non-malignant microenvironment cell types across the 7 unique tumors of the cohort (Supplementary Fig. 8A-B). We also modelled cell-cell signaling events that occur in PNTs, identifying possible cell extrinsic regulatory mechanisms within the tumor milieu. Our samples showed significant cellular heterogeneity and we identified rare subpopulations in a number of samples in this cohort, which would have otherwise been masked in bulk gene expression discovery. This highlights the utility of scRNA-seq in the classification of cell diversity that exists within heterogenous tumors.

Using single-cell sequencing, we could facilitate virtual microdissection of malignant cells and microenvironmental heterogeneity within and between tumors. In particular, among malignant populations, we detected the expression of genes related to mesenchymal/neural crest cell and adrenergic/noradrenergic neuroblastoma cells, consistent with previous reports^7,8,36,37^. Notably however, we found that some high-risk tumors display an additional transitional state with gene expression characteristics associated with aggressive disease, rapid proliferation and metastasis. Consistent with these findings, transitional signatures were also associated with more aggressive clinical behavior in bulk transcriptome cohorts, such as enrichment in high-stage and MYCN-amplified patients and a strong association with poor prognosis. This enrichment with MYCN-amplification was particularly intriguing since none of the tumors of this cohort were MYCN-amplified, more evidence that high-risk neuroblastoma is often a MYC-driven disease regardless of whether the tumor is MYCN-amplified or MYCN non-amplified^38-40^. Moreover, while only based on the small sample size of this cohort, the 2 tumors which had relatively high proportions of transitional cells (NB3 & GNB1), were the only tumors that failed to show a complete response to upfront therapy (Supplementary Table 1), again highlighting the potential importance of transitional cells as biomarkers of prognosis and target for novel therapies in PNTs.

Prior studies have speculated that adrenergic/noradrenergic cells transdifferentiate to mesenchymal/neural crest cells and vice versa^7,8,36,37,41^. This theory was primarily based on spontaneous state polarization of isogenic pairs of cells and forced overexpression experiments using strong lineage associated transcription factors to reprogram cell phenotype such as PHOX2B, PRRX1, ASCL1 and NOTCH3^7,8,36,41^. Our trajectory modelling using single-cell RNA-seq data, and our reclassification approach for H3K27ac landscapes in neuroblastoma cell lines is in support of this theory, albeit by a slightly more complex mechanism involving transitional cells as a transdifferentiation intermediate cell state. This model is akin to the long-held observation that neuroblastoma cell lines demonstrate 3 morphological variants: neuroblastic (N), substrate-adherent (S) or intermediate (I) phenotypes^42-45^. N-type cells are said to resemble embryonic sympathoblasts (similar to the adrenergic phenotype), S-type cells resemble Schwannian, glial or melanocytic progenitors (similar to the mesenchymal phenotype) and I-type cells have an intermediate phenotype with the potential to differentiate toward either cell type (similar to our proposed transitional phenotype)^42-45^. Importantly, similar to the 3-state model we present here, neuroblastoma cells in culture are capable of interconversion and transdifferentiation between these morphological classes^42-45^. Moreover, examination of scRNA-seq data derived from the embryonic murine sympathoadrenal system, shows expression patterns that suggest that differentiation through a transitional state may be developmentally co-opted. This phenotype has distinct gene expression markers of significant note such as MYCN and EZH2, as well as upregulation of neurodevelopment and neurogenesis-related pathways and activation of some distinct super-enhancer loci. We suggest that the transitional state is likely transient in the respect that the cells have greater cell plasticity and are more likely to adapt to changing environments. We hypothesize that this state could act as a pit-stop during transdifferentiation depending on subtle environmental exposures, which may also give them a fitness advantage to survive anti-cancer barriers (i.e. anti-cancer therapies, and/or homeostatic anti-cancer mechanisms). Nevertheless, future research will benefit from functional genomics studies on key “transitional” genes and lineage tracing studies that can follow phenotypical adaptation in response to environmental stimuli. This will be required to elucidate the precise interconnections between all cell states, and how these states relate to other established models of neuroblastoma cell phenotype such as morphological and epigenetic classification methods.

Another key finding from recent studies, is that tumor cells convert to a mesenchymal/neural crest cell phenotype in relapsed neuroblastoma tumors and upon drug exposure *in vitro*, suggesting that mesenchymal differentiation is a drug-resistance mechanism in neuroblastoma^7,8,46^. However, our analysis of bulk gene expression from diagnostic tumors paradoxically shows that the mesenchymal-only class is not a predictor of poor outcome, and indeed is generally predictive of favorable outcome. Since our data showed that transitional signatures are more predictive of poor outcome, we suggest that a mesenchymal phenotype is not a high-risk phenotype per se, but rather that transitional cells may have the capacity to escape cytotoxic therapy by transdifferentiation to mesenchymal phenotype under drug selection pressure. But importantly, since cells maintain greater plasticity, this would potentially also allow adaptation back a cell state that supports other characteristics of aggressive tumors when drug-selection pressure is released (e.g. rapid proliferation and metastasis). This concept is supported by the analysis of transitional subgroups in bulk neuroblastoma cohorts, which demonstrates transitional-only and transitional-mesenchymal subgroups have significantly poorer outcome than other classes, including transitional-adrenergic class. This suggests that a mesenchymal differentiation tendency is actually unfavorable rather than an intrinsic mesenchymal phenotype.

Among non-malignant cells, we found that non-neuroblastoma samples have a higher proportion of infiltrating immune cells compared with neuroblastoma samples, especially cytotoxic T-cells. Cytotoxic T-cells are a key player in tumor surveillance and inhibition of tumor growth by secreting cytokines as well as perforin or Fas-mediated cytotoxic response^47^. Therefore, it is an important target of immunotherapy and considered as an important prognostic indicator in many cancers, including neuroblastoma^29,48^. Multiple studies have shown that T lymphocyte infiltration largely correlates with tumor differentiation status and *MYCN-*non-amplified gene status^29^. MYCN has been reported to downregulate the expression of peptide-MHC I (pMHC I) in neuroblastoma to evade host immune surveillance leading to a T-cell-poor microenvironment in MYCN-amplified neuroblastoma^49^. While our cohort lacked MYCN-amplified samples, it was interesting to note that MYCN upregulation in transitional neuroblasts may induce similar changes in the high-risk samples identified to contain these cell types.

Using modelling of cell-cell signaling in our cohort, an immune-regulatory relationship between cytotoxic T-cells and malignant cells was also inferred, with the cytotoxic T-cell specific ligand ITGB7 predicted to effect the adrenergic gene target ZFHX3 by binding to integrin or hedgehog signaling molecules on neuroblasts (ITGAE, ITGA9, PTCH1, PTCH2). ZFHX3 is a transcription factor involved in normal neurogenesis/neuronal differentiation and is highly expressed in favorable NB cases. Furthermore, the upstream integrin/hedgehog signaling molecules had also been implicated in NB differentiation suggesting that this interaction with cytotoxic T-cells may have eventuated in a neuronal differentiation phenotype in neuroblasts^50-53^. This finding supports the aforementioned connection between the differentiation status of neuroblasts and the extent of T cell infiltration in NB tumors. Interestingly, we also found a relatively high level of macrophages in neuroblastoma tumor microenvironments compared with ganglioneuroblastoma and ganglioneuroma samples. From our cell-cell interaction modelling we had also predicted that IGF1 ligands released specifically by non-inflammatory macrophages could modulate the transitional gene target MYCN by binding to IGF1R receptors on neuroblasts. MYCN is a core pro-tumorigenic transcription factor in malignant neuroblasts and is strongly associated with poor patient prognosis, the upstream IGF1R receptor is highly expressed in neuroblastoma and is synonymous with a metastatic phenotype, suggesting an increased metastatic potential in neuroblasts after IGF1R mediated interactions with non-inflammatory macrophages^54-58^. This metastasis promoting interaction is further supported as tumor-associated macrophages are known to be more prevalent in neuroblastoma patients with metastatic disease compared with localized lesions^13^. In line with these findings we had also predicted that AREG ligands released by inflammatory macrophages would influence the mesenchymal gene target NUPR1 by interacting with EGFR family receptors (EGFR, ERBB2, and ERBB3) on malignant neuroblasts. NUPR1 is a transcriptional regulator that has not previously been implicated in neuroblastoma but has been established in other cancers as a metastasis and drug-resistance promoting gene, the upstream EGFR receptors have, similarly, been associated with a metastatic phenotype in neuroblastoma, supporting the notion that EGFR mediated interactions with inflammatory macrophages elicit a metastatic phenotype^59-62^. The identification of non-malignant cell types and potential cell-cell interactions have provided a glimpse into the complex paracrine signaling events that may underlie malignant neuroblast transdifferentiation in individual PNTs, though there still remains ample room for experimentally validating these interactions using cell line models or spatial transcriptomic methodologies in PNT tumors.

Although scRNA-seq provides new tools for high resolution profiling of cell populations, it is still with some limitations. One limitation of our study is that the high-risk primary tumors were obtained from patients who had received prior chemotherapy, while the remaining tumors in the study were chemotherapy naïve. Therefore, it is unknown to what extent these treatments may have influenced intratumoral heterogeneity. This may be particularly important when considering immune proportions, because of the potential immuno-depletive effects of chemotherapy. Future longitudinal studies that pair pre-treatment biopsy specimens with post-chemotherapy surgical resections and relapsed tumors will be valuable to unveil tumor phenotype adaptation in the context of drug induced tumor evolution. Another limitation within our study is the small number of samples profiled. This is caused by the relative paucity of ganglioneuroblastoma intermixed (GNBi) and ganglioneuroma, as well as the difficulty of obtaining adequate viability to create viable cDNA libraries after single-cell dissociation and cell sorting. The addition of more samples with a diverse molecular background will be invaluable to the single-cell field such as MYCN-amplified, ALKmut, ATRXmut, PHOX2Bmut, TERT rearrangements, RASmut, p53mut and other segmental chromosomal alterations recurrent to neuroblastoma^63,64^. This will facilitate the generation of molecular-associated single-cell gene expression profiles that will be important to distinguish genetic and non-genetic facets of cell phenotype. Finally, while we have aimed to assess tumor cellular compositions in a quantitative manner, it will be important to validate these findings with potential that there was a bias in cell collection that arises when enriching viable cells. A notable cell type that was not identified was the Schwann cells when there was histological evidence of Schwannian stroma, highlighting the potential for missing or inaccurately quantifying cell types in the tumor microenvironment.

In conclusion, our analysis has uncovered extensive intra- and inter-individual, functional and transcriptomic heterogeneity in neuroblastic tumor cells and associated microenvironmental cells. We identify novel mechanisms of phenotypical heterogeneity, highlighting the importance of considering cell plasticity and transdifferentiation potential in new therapeutic strategies for PNTs.

## Methods

### Patient sample collection

Fresh peripheral neuroblastic tumor (PNT) tissue samples were obtained from 7 patients enrolled at the Xinhua Hospital affiliated to Shanghai Jiao Tong University School of Medicine (Shanghai, China). This study was approved by the ethics committee of Xinhua Hospital and was conducted in accordance with the Declaration of Helsinki. Patients were enrolled for this study based on pathological diagnosis. All patients were enrolled under informed consent. Staging and risk assessment were performed according to the International Neuroblastoma Staging System Committee (INSS) system and the International Neuroblastoma Risk Group (INRG) respectively, based on tumor histology and MYCN-fluorescent in situ hybridization (FISH) analysis. Tumors were resected from the primary site either prior to the patient undergoing chemotherapy or post-induction chemotherapy. Treatment regimen included “treatment protocol CCCG-NB-2014”. Detailed clinical information on individual patients in this study are outlined in Supplementary Table 1.

### Immunohistochemical staining

All PNTs specimens were fixed, paraffin-embedded, sectioned, and stained with hematoxylin and eosin (H&E) following routine method of Xinhua Hospital’s Pathology. Immunohistochemical (IHC) studies employed 5μm paraffin-embedded slides. Antigen was retrieved by citric acid buffer (PH6.0) in the microwave oven. Endogenous peroxidase was inactivated by incubation in 3% H2O2 for 25 minutes. After using 3%BSA to block nonspecific sites for 30 minutes, slides were incubated with primary antibody anti-CD3 (1:100, GB13014, Servicebio) to assess the presence of infiltrating T lymphocytes.

### Single cell preparation and flow cytometry

Following surgical resection, fresh tissue samples were immediately transferred to DMEM/F12 medium (MULTICELL), supplemented with 2% FBS (Gemini Bio-Products) and were delivered on ice within 60 minutes to the laboratory to be processed. Tissue processing was completed within 90 minutes of collection. Samples were first rinsed in ice cold PBS (MULTICELL) with 2% penicillin-streptomycin (BBI life science) and then minced into 1mm3 pieces using curved scissors under sterile conditions. The fragments were further enzymatically dissociated into single cells using LiberaseTM (Roche) with a final concentration of 0.26 U/mL and a final volume of 25ml Liberase solution/0.5g tissue on a shaker with a speed of 85 rpm for 45 min at 37°C. The Liberase was diluted in Hyclone L-15 Leibovitz media. DMEM/F12 media supplemented with 10% FBS was used to stop dissociation. The resulting single-cell suspension was filtered through a 70µm nylon cell strainer (BD Falcon) and then centrifuged for 5 min at 300g at room temperature. The cell pellet was resuspended in 500µL of PBS supplemented with 0.1% BSA (BBI life science) and passed through a 40µm nylon cell strainer (BIOFIL). This cell suspension was stained with Calcein-Blue (Invitrogen) and DRAQ5 (CST) prior to fluorescence activated cell sorting (FACS) in order to isolate live and nucleated cells (Calcein-Blue+ DRAQ+). Single cells were sorted using a BD Becton Dickinson FACSAriaII into 96-well PCR plates. Each well of the 96-well plate contained 3µL lysis buffer (10U Recombinant RNase Inhibitor (Takara Bio), 0.2% Triton X-100 (Sigma-Aldrich), 3mM dNTP mix (Takara Bio)), and reverse transcription reagents (see below). Immediately following FACS, plates were briefly centrifuged and stored at −80°C.

### Single cell library preparation and RNA sequencing

scRNA-seq was conducted using the Smart-seq2 protocol^9^ with some modifications being made to incorporate unique molecular identifiers (UMIs) into the 3’ end of transcripts^65^. Reverse transcription was performed directly in the 96-well plates by incubation at 72°C for 5 min, after which the plate was replaced on ice to allow the oligo-DT primer to hybridize to the poly(A) tail of the mRNA molecules. The oligo(dT) primer used in reverse transcription included an additional 8bp cell barcode, 9bp unique molecular identifiers (UMIs) and template-switching oligo sequence. PCR amplification was performed by adding 15µL PCR mix containing 0.5U KAPA HiFi HotStart (Kapa Biosystems), 1x KAPA Buffer (Kapa Biosystems), 12.5mM MgCl2 (Kapa Biosystems), 5µM ISPCR Primer and 7.5mM dNTP mix (Takara Bio). PCR amplification was performed in a thermal cycler (BIORAD C1000 Touch Thermal Cycler) at 98°C for 3 min, 24 cycles of 98°C for 20s, 67°C for 15s, and 72°C for 6 min, and a final incubation at 72°C for 5 min. PCR products were purified using 1X AMPure XP bead (Beckman, Cat. #A63882) and Qubit dsDNA HS Assay Kit (Invitrogen, Cat. #Q32854). TN5 tagmentation and library amplification realized 3’end fragments by Nextera XT DNA Sample Preparation Kit (Illumina, Cat#FC-131-1024) according to the manufacturer’s instructions while P5_TSO and Nextera_N7xx took the place of the custom primers. Pooled single cell libraries were sequenced at an average depth of 0.5 million reads per cell on an Illumina HiSeqX10 instrument using a 2 x 150bp paired end sequencing kit (Genewiz).

### Single cell RNA-seq data analysis

Raw sequencing data was processed using the Dr.Seq2 pipeline^66^. Reads were aligned to the human genome (hg19) using STAR^67^. In each read, cell barcodes were between 9:16bp and UMIs were between 17:26bp. Following the Dr.Seq2 pipeline we converted the resultant aligned sam file to a bed file using a custom script. We then generated a gene annotation file and annotated the aligned bed file. We then reproduced the aligned sam file to contain gene annotations, cell barcode and UMI information for each read. UMI counts were calculated by removing duplicate reads which had an identical genomic location, cell barcode and UMI sequence, resulting in the final UMI count matrices used for downstream analyses. Cells with fewer than 600 genes or more than 5000 genes were removed (genes were only considered if they were expressed (UMI > 1) in at least 3 cells). Furthermore, cells with more than 30% ribosomal gene content were filtered. Downstream log normalization and scaling were performed using the R package Seurat (version 3)^68^. Highly variable genes were identified using “vst” model of all expressed genes in Seurat. This output was used to perform a principal component analysis (PCA), and the variance in each principal component (standard deviation) was visualized on an elbow plot which further used to determine the PC-cutoff for downstream clustering and dimensionality reduction analysis. Additionally, clustering trees were generated using the R Clustree Package^69^ to identify a stable clustering resolution parameter. After identifying the number of PCs and resolution to use for downstream analysis, Uniform Manifold Approximation and Projection (UMAP)-based dimensional reduction was performed using Seurat^68^. All single-cell expression signatures were created using AddModuleScore function in Seurat and visualization of data was undertaken using ggplot2, Seurat, Monocle2, and CONICSmat plotting functions^16,19,68,70^.

Copy number variations were inferred from single cell RNA-seq data using the CONICSmat R-package^16^ with some minor modifications. Rather than using chromosomal arms for regions of mixture model assessment, a custom script was used to define a regions of interest. This script defined regions based on the difference between the 100 gene rolling average of inferred malignant and non-malignant cells (for this, “sympathetic” and “mesenchymal” clusters were assumed to be malignant, while the non-malignant clusters were assumed to be all other clusters). Statistically significant mixture-models were then assessed using these new differentially defined regions. All reported mixture models were chosen based on: a 2-component model being more likely than a 1 component model using Bayesian Information Criterion (BIC) ratio of >1.01 (see Supplementary Table 2, BIC ratio = BIC.1component/BIC.2.component). After candidate regions for CNVs were chosen, a CNV score was calculated for each cell as the average adjusted area under the curve (adj.auc) for all candidate regions identified in that sample, using the 100 gene rolling average difference between malignant and non-malignant cells (the auc was adjusted for the length of each region in terms of number of genes). All CNV heatmaps were generated using the identified malignant and non-malignant clusters from the above analyses and were plotted using an expression threshold to keep only genes with average expression greater than the 10% quantile of average gene expression for all genes. For visualization, CNVs were color mapped using a lower bound of 1 standard deviation and upper bound of 2.5 standard deviations from the mean of the global rolling average. Projection of CNVs to UMAP plots was done using the Seurat^68^.

Trajectory analyses of single cells was performed using the Monocle2 R package (^19^ version 2.8.0) with the CellDataSet created using UMI count values. Pseudotime and trajectory calculation was undertaken using expressed genes from the Adrenergic and Mesenchymal signatures previously published^8^. Gene expression pseudotime plots were plotted using plot_genes_in_pseudotime function and gene expression pseudotime heatmaps were plotted using plot_genes_branched_heatmap function in the Monocle package. Cell state signatures were created using the top 60 genes ranked by adjusted p-value using FindAllMarkers for overexpressed genes (average log fold change>0.25) that are expressed in at least 25% of cells for that state designation (Supplementary Table 3).

Gene ontology analysis was performed using the topGO package. Significantly enriched gene ontology terms were identified using the “classic” algorithm and the “fisher” test. Representative tests were chosen that had a p-value < 0.001.

Ligand-receptor-target interaction modelling in single cells was performed using the NicheNet R package^30^. The approach included a number of steps to identify higher confident interactions, as summarized in Supplementary Fig. 7A. In order to exclude lowly expressed/represented ligands and receptors in our analysis we applied gene expression and cell proportion based cut-offs around the non-zero mean (Supplementary Fig. 7B). We considered that each patient had a unique tumor microenvironment and so ran NicheNet models on a per-patient basis, incorporating malignant and non-malignant cell types that were only present in that patient (≥5 cells) (Supplementary Fig. 7C). Differentially expressed genes in adrenergic, transitional and mesenchymal neuroblasts were modelled as downstream gene targets of ligands produced by non-malignant cell types in each patient. Ligand activity scores were generated for each malignant cell and further correlated with target gene/signature expression using Pearson regression to better infer whether a given ligand effected target gene/signature expression. These interactions had to be both positively and significantly correlated (p<0.05 & r>0) to be considered for downstream analysis (Supplementary Fig. 7D). We finally looked for overlapping interactions among patients (≥2 patients) which signified ligand-receptor-target interactions that were likely to be more representative. Overlapping ligand-target interactions were visualized as chord diagrams, with chords representing interactions between a specific ligand in one non-malignant cell type and one/several downstream gene targets in neuroblasts (Supplementary Fig. 7E). Ligand-target and ligand-receptor scores from NicheNet were visualized alongside average expression values of ligand/receptors/targets in single cells to further illustrate interaction networks for overlapping interactions (Fig.7A).

### Bulk Tumor Analyses

Bulk tumor analyses were conducted on previously described tumor microarray cohorts: Kocak^26^ and Versteeg^27^. Average gene expression signatures (i.e. Adrenergic, Transitional and Mesenchymal) were created from the average of all z-scores for genes in each signature. Kaplan-Meier analyses were conducted based on a median expression cutoff to stratify high and low expression groups. P-values for survival analysis were calculated using log-rank tests. To classify tumors into 5 “Tumor Class” subgroups, average z-scores for each of the Adrenergic, Transitional and Mesenchymal signatures were again z-score scaled and subgroups were created based on the following criteria:

| Adrenergic signature expression | Transitional signature expression | Mesenchymal signature expression | Assignment |
| --- | --- | --- | --- |
| >0.35 | <0.35 | <0.35 | Adrenergic |
| >0.35 | >0.35 | <0.35 | Adrenergic-transitional |
| <0.35 | >0.35 | <0.35 | Transitional |
| <0.35 | >0.35 | >0.35 | Transitional-Mesenchymal |
| <0.35 | <0.35 | >0.35 | Mesenchymal |
| <0.35 | <0.35 | <0.35 | Either Adrenergic, Transitional or Mesenchymal groups, based on which signature had the highest value |
| >0.35 | <0.35 | >0.35 | Either Adrenergic or Mesenchymal groups, based on which signature had the highest value |
| >0.35 | >0.35 | >0.35 | Either Adrenergic-transitional or Transitional-mesenchymal groups, based on whether the Adrenergic or Mesenchymal signature had the highest value |

### H3k27ac ChIP-seq & RNA-seq data integration and analysis

bigWig files containing H3k27ac ChIP-seq read densities for 33 neuroblastoma samples were retrieved directly from the Gene Expression Omnibus (GEO); GSE90683^7^. bigWig files were converted to the bedGraph format using the UCSC *bigWigToBedGraph* tool, and then combined into one bedGraph using bedtools (*unionbed*)^71^. Each sample was then normalized for total read density and multiplied by a 10^6 scaling factor. The bedops tool was then used to find overlapping read bins with the 5975 previously annotated superenhancer regions^7,72^. Matched RNA-seq profiles for these 33 samples were directly retrieved from GEO; GSE90683^7^. Average gene expression signatures (i.e. Adrenergic, Transitional and Mesenchymal) were created from the average of all z-scores for genes in each signature. To classify samples into the same 5 subgroups as described above for bulk tumor analyses, average z-scores for each of the Adrenergic, Transitional and Mesenchymal signatures were again z-score scaled and subgroups were created based on the criteria described previously. Normalized ChIP-seq read densities in the bedGraph file were then averaged across matching genomic bins for samples belonging to each of the five subgroups determined by RNA-seq. Five bigWigs representing averaged H3k27ac ChIP-seq profiles across the five subgroups were generated from the bedGraph file using bedtools (*bdg2bw*)^71^. Differentially enriched super-enhancers were then identified using 2-sided t-tests for the normalized ChIP-seq read densities for previously annotated super-enhancer regions^7^ between cells belonging to each Tumor Class. All p-values were then adjusted using Benjamini & Hochberg correction and enriched super-enhancers were identified if p<0.05. Enriched super-enhancers for the adrenergic cell lines were identified by comparing to the mesenchymal and vice versa. Transitional enriched super-enhancers were identified by comparing cell lines of any combination of Adrenergic-Transitional, Transitional or Transitional-mesenchymal cell lines compared with Adrenergic & Mesenchymal cell lines (see Supplementary Table 4A-B for comparisons and statistics). Downstream visualization was undertaken using the EnrichedHeatmap package on all unique enriched super-enhancers identified by differential testing based on 1kb bins through the 0.5Mb region around the center of each annotated super-enhancer locus^73^. All H3K27ac z-score plots were created by taking the mean of all z-scores of all enriched superenhancers designated as either Adrenergic, Transitional or Mesenchymal for each binned region for each cell line and plotted using ggplot2^70^. H3K27ac z-score plots for Adrenergic and Mesenchymal enriched super-enhancers were then quantified for each tumor class by calculating the total area under curve using the *auc* function in R package MESS using default parameters considering each cell line belonging to the 5 Tumor Classes (error bars among cell lines from each class are represented as standard deviation). H3K27ac traces on core-regulatory transcription factors and markers of transitional state were plotted using the Gviz package^74^ using regions corresponding previously annotated super-enhancers^7^.

## Supplementary Figure Legends

**Supplementary Figure 1**

A) Schematic representation of the sample preparation workflow for generating scRNA-seq profiles from primary peripheral neuroblastic tumors. Surgically resected tumors were first enzymatically dissociated into a single cell suspension. Viable and nucleated cells were then sorted into 96-well plates prior to a modified Smart-seq2-based library preparation and next generation sequencing.

B) Fluorescence-activated cell sorting (FACS) was performed on dissociated tumor samples to isolate viable and nucleated single cells. Representative figure showing the gating strategy after staining cells with DRAQ5 (nucleated cells) and Calcein Blue (viable cells).

C) Flowchart depicting the steps involved in data pre-processing of demultiplexed .fastq using the DrSeq2 pipeline and filtering/post-processing with the SeuratV3 (R 3.5).

D-F) Histograms showing the distribution of total (D) UMI counts, (E) unique genes and (F) ribosomal reads expressed per cell. Histograms are color coded according to original patient identity. Red dashed lines indicate cutoffs used for selecting high quality cells.

G) Principle component analysis (PCA) elbow plot, representing the standard deviation of each principal component, was constructed to visualize the amount of variance in each PC. Here, the plot shows an elbow at PC 10, which was chosen for PC cut-off for downstream data analysis, namely clustering and dimensional reduction.

H) Clustering trees based on increased resolution parameters of the Louvain clustering algorithm used in Seurat. This plot was generated using the Clustree package (R 3.5)^69^. A stable clustering resolution parameter of 0.3 was chosen for downstream analysis.

I) Gene expression heatmap of the top differentially expressed genes (rows) across each transcriptomic cluster (annotated columns). Individual columns represent cells, and a scale bar is provided indicating gene expression z-scores.

J) Violin plot of average gene expression levels of markers that define broad cell types colored according to clusters (Fig. 1D). Points within the violin represent individual cells

K) UMAP projection of CNV score in all cells (n=4267) from the 7 patient samples.

L) Heatmap of CNVs for individual cells (rows), visualized by a rolling gene expression average centered on 100 gene windows at each chromosomal position (columns). Dark-red: copy number gains, blue: copy number losses. Colored sidebars denote cell type. Malignant cells (above dashed line) display an aberrant copy number profile compared to non-malignant cells (below dashed line).

**Supplementary Figure 2**

A) UMAP plots of 2024 malignant cells from all 7 tumor samples. Cells are colored by transcriptomic cluster identity.

B) Gene expression heatmap of the top differentially expressed genes (rows) genes in each malignant cell cluster (annotated columns) that were defined in Supplementary Fig. 2A. Individual columns represent cells and a scale bar is provided indicating gene expression z-scores.

C) UMAP of 2024 malignant cells colored based on the average expression of genes in a previously published adrenergic gene signature^7^ (blue: low, dark-red: high).

D) UMAP of 2024 malignant cells colored based on the average expression of genes in a previously published mesenchymal gene signature^7^ (blue: low, dark-red: high).

E) Trajectory plot of all malignant cells (n=2024) constructed by Monocle 2 (DDTRee). Points represent cells and are colored by original sample identity.

F) Proportion of malignant cells that are either in an; adrenergic, transitional or mesenchymal cell state, within each tumor sample.

**Supplementary Figure 3**

A) UMAP plots of all malignant cells (n=2024) from the 7 tumor samples. Dataset is colored according to the cell state generated using Monocle DDRTree.

B) Gene expression heatmap of the top 60 differentially expressed genes (rows) of cells in either an; adrenergic, transitional or mesenchymal cell state (annotated columns). Individual columns represent cells and a scale bar is provided indicating gene expression z-scores. These genes represent the top 60 genes in the Adrenergic, Transitional and Mesenchymal gene signatures used in subsequent analyses.

C) Heatmap of gene expression signatures ordered by pseudotime along arms of the trajectory in Fig. 2G. Pseudotime was calculated considering the adrenergic to transitional trajectory (left column) and the transitional to mesenchymal trajectory (right column). Sidebars annotate either the cell type (columns) or the gene signature type (rows). Scale indicates gene expression z-scores for the given trajectory.

D) Gene ontology (GO) enrichment analysis of adrenergic, transitional and mesenchymal gene signatures. Representative GO terms that were significantly enriched in each gene set are shown.

E) Trajectory plot of all malignant cells colored based on the average expression of genes within the GO gene-sets identified in Supplementary Fig. 3D. Scales indicate average gene expression of all genes in the signature (blue: low, dark-red: high expression).

F) Violin plots showing the expression of the adrenergic, transitional and mesenchymal state signatures in murine neural-crest cells from embryonic day 12.5. Cells are grouped by the cell types as defined in Furlan *et al*^24^. SCPs; Schwann-cell precursors.

G) Proportion of actively dividing or non-dividing cells in adrenergic, transitional and mesenchymal cells. Cycling states were annotated based on the average expression levels of genes within the S and G2M (dividing) and G1 (non-dividing) gene sets previously published^17,18^.

H) Violin plot illustrating significant difference in the disseminated tumor cell (DTC) score between adrenergic, mesenchymal or transitional cells. A disseminated tumor cell score was determined for each cell group by determining the difference between the average expression of significantly up-regulated and down-regulated genes of a previously published DTC gene set^25^. P-values are reported from a t-test. Points represent individual cells.

**Supplementary Figure 4**

A) Kaplan-Meier plots of overall survival (top) and event free survival (bottom) in 649 neuroblastoma patients from the previously published Kocak cohort^26^ based on the expression levels of genes in the Adrenergic, Transitional or Mesenchymal gene sets. Patients were dichotomized by the median expression of these gene signatures (n=324 & 325 tumors respectively). P-values determined using log-rank tests comparing both groups.

B) Kaplan-Meier plots of overall survival (top) and event free survival (bottom) in 88 neuroblastoma patients from the previously published Versteeg cohort^27^ based on the expression levels of genes in the Adrenergic, Transitional or Mesenchymal gene sets. Patients were dichotomized by the median expression of these gene signatures (n=44 & 44 tumors respectively). P-values determined using log-rank tests comparing both groups.

C) Heatmap detailing the classification of neuroblastoma patients from a bulk mRNA (microarray) profiling cohort^27^ based on adrenergic, transitional and mesenchymal gene signatures. Gene expression values (rows) for each patient (columns) were arranged based on their respective gene signature. Column averages were used for each signature (shown immediately below the heatmap) to classify patient to Tumor Class subgroups: Adrenergic-only, Adrenergic-Transitional, Transitional-only, Transitional-Mesenchymal, and Mesenchymal-only (see methods). Patient metadata concerning; *MYCN* status (amplified) and neuroblastoma staging (INSS 4) are annotated in bars underneath the heatmap. Scale bar indicates gene expression z-scores.

D) Kaplan-Meier plots of overall survival (top) and event free survival (bottom) in the Versteeg cohort based on Tumor Class classifications made in Supplementary Fig. 4C.

E) Boxplots of transitional gene signature expression in patients of the Versteeg cohort (n=88), when dichotomized by MYCN amplification status (left; non-amplified vs amplified) or INSS staging (right; stage 1, 2, 3 & 4S vs stage 4). P-values are reported from 2-sided t-tests, and boxes represent first-quartile/median/third-quartile of expression values, with the whiskers representing the 95% confidence interval, and large points representing outliers. Small points represent individual tumors.

**Supplementary Figure 5**

A) Heatmap detailing the classification of neuroblastoma cell lines from a bulk mRNA (RNA-seq) profiling cohort (n=33 cell lines^7^) based on adrenergic, transitional and mesenchymal gene signatures. Gene expression values (rows) for each cell line (columns) were arranged based on their respective gene signature. Column averages were used for each signature (shown immediately below the heatmap) to classify cell lines to Tumor Class subgroups: Adrenergic-only, Adrenergic-Transitional, Transitional-only, Transitional-Mesenchymal, and Mesenchymal-only (see methods). Cell line metadata concerning; *MYCN* status (amplified), ALK mutation status, PHOX2B mutation status and prior super-enhancer classification (SE Type) from Boeva et al^7^ are annotated in bars underneath the heatmap. Scale bar indicates gene expression z-scores.

B) Global normalized H3K27-acetylation (H3K27ac) values for all 33 cell lines. Super-enhancer loci enriched in Adrenergic (top) or Mesenchymal (bottom) subgroups are shown. H3K27ac values are representative of each cell line and represent 0.5Mb centered on the super-enhancer locus in that region. Bin size for each trace is 1kb.

C) Global normalized H3K27ac values for adrenergic superenhancers (red) and mesenchymal super enhancers (blue) were quantified for each of the 33 cell lines using a z-score value of the total area under the H3K27ac curve.

D) Global normalized H3K27-acetylation (H3K27ac) values for all 33 cell lines (top) or when classified to 5 subgroups (bottom). Super-enhancer loci enriched in Transitional cell lines are shown (n=94). H3K27ac values are representative of each cell (top) or averaged across all cell lines classified in that group (bottom) and represent 0.5Mb centered on the super-enhancer locus in that region. Bin size for each trace is 1kb.

E) Representative H3K27ac tracks for each group of classified cell lines for super-enhancer loci shown to be enriched in Transitional-grouped cell lines. Dark-green box is an example of Adrenergic-Transitional super-enhancer locus, Green boxes are examples of Transitional or pan-transitional super-enhancer loci, Light-green box is an example of a Transitional-Mesenchymal super-enhancer locus.

**Supplementary Figure 6**

A) Gene expression heatmap of the top differentially expressed genes (rows) across non-malignant cell clusters (annotated columns) that were defined in Fig.6B. Individual columns represent cells and a scale bar is provided indicating gene expression z-scores.

B) UMAP plot of 2243 non-malignant cells from 7 tumors with cells colored based on the average expression of marker genes for specific cell types (marker genes and associated cell types are indicated above to each plot). Scale: average gene expression z-scores (blue: low, dark-red: high). Left: Ellipses indicate the initial cell type classification. Right: Arrows indicate regions of high expression for specific cell subtype classification.

C) Frequency of T-cell subtypes (top), macrophage subtypes (bottom) within each tumor sample as well as in the major tumor types (neuroblastoma compared with ganglioneuroblastoma/ ganglioneuroma).

D) Representative immunohistochemical staining of CD3 in the 7 tumor samples indicating the presence of T cells. 40X objective (equivalent to 400X magnification). Scale bar: 20μm.

**Supplementary Figure 7**

A) Workflow for NicheNet modelling describing steps taken to determine overlapping ligand-receptor-target interactions among the 7 PNT patients.

B) Upper panels display frequency histograms of ligand and receptor expression (normalized expression values) in single cells, red lines indicate the non-zero mean of all values which is also provided in each plot. Lower panels display frequency histograms of ligand and receptor proportions among single cells (>0 normalized expression value in a given cell), red lines indicate the non-zero mean of all values which is also provided in each plot.

C) Heatmap of cell type inclusion criterion for the NicheNet model. On a per-patient basis, only cell types that had more than 5 cells (black) were considered for NicheNet modelling (n=61). Absent cell types (or those that only consisted of less than 5 cells) (white) were not considered for NicheNet modelling. Columns represent non-malignant cell types whilst rows represent malignant cell types in each patient. Some models did not produce any significant interactions between associated cell types (n=7). The heatmap to the right represents those cell types which passed the threshold (≥5 cells) in each patient.

D) Scatter plot of Pearson regression coefficients (r) of correlated ligand activity scores in malignant cells against target gene (x-axis) or target signature (y-axis) expression. Significant (p<0.05) and positive correlations (r>0) are colored in red (n=961) and their density are summarized by contour lines.

E) Representative chord diagram of predicted ligand-target interactions (n=87) between non-malignant cell types (bottom) and malignant neuroblasts (top) using NicheNet. Ligands shown (n=24) are expressed in non-malignant cells. Individual chords are drawn from the expressed ligand (n=24) to the predicted gene target (n=30) in neuroblast subsets. Individual chords are colored by cell type specificity of the ligand and are annotated in the figure. All ligand-target links shown were predicted independently in at least 2 patients.

F) Stacked column graphs showing the proportion of ligand-receptor-target links (n=87) between adrenergic, transitional and mesenchymal neuroblasts and non-malignant cells. Bars are colored by non-malignant cell type specificity; annotations are displayed to the right.

**Supplementary Figure 8: Summary of PNT single-cell RNA-sequencing**

A) Left: UMAP plot showing all cells sub-classified into various subtypes, Right: the proportion of non-cell types within each tumor and major tumor types are also shown (neuroblastoma compared with ganglioneuroblastoma/ganglioneuroma). Colors indicate the various subtypes.

B) Summary table of the various samples used in this study.

## Supplementary Table Legends

**Supplementary Table 1**

Summary of relevant patient information, from whom primary site tumors were isolated and profiled by scRNA-seq.

**Supplementary Table 2**

Genomic regions with chosen 2-component Gaussian mixture models and accompanying Bayesian Information Criterion (BIC) statistics

**Supplementary Table 3**

Gene signatures for each malignant cell state. Top 60 genes identified by Seurat function FindAllMarkers.

**Supplementary Table 4**

Differential statistics for enriched super-enhancer regions related to Figure 5.

**Supplementary Table 5**

NicheNet summary statistics for significant cell-cell signaling interactions related to Figure 7.

## Acknowledgments

The work was supported by grants from the National Key Research and Development Program of China (Grant No. 2017YFA0103902); The National Natural Science Foundation of China (Grant No. 81570760 and 31771283); Project of Shanghai Committee of Science and Technology, China (Grant No. 16411962500); Translational Medicine Innovation Fund of Shanghai Jiao Tong University School of Medicine, China (Grant No. 15ZH2005); the Fundamental Research Funds for the Central Universities of Tongji University. W.S. was supported grants from National Key Research and Development Program of China (2018YFD0900600), The National Natural Science Foundation of China (41676119) and the Fundamental Research Funds for the Central Universities from the Ocean University of China. D.C received Cancer Institute NSW Early Career Fellowship #DC000594.

## Compliance with ethics guidelines

The authors declare no conflict of interest.

