## Supplementary Figures for "Single-cell RNA-sequencing of peripheral neuroblastic tumors reveals an aggressive transitional cell state at the junction of an adrenergic-mesenchymal transdifferentiation trajectory"

Supplementary Figure 1 (1 of 3)

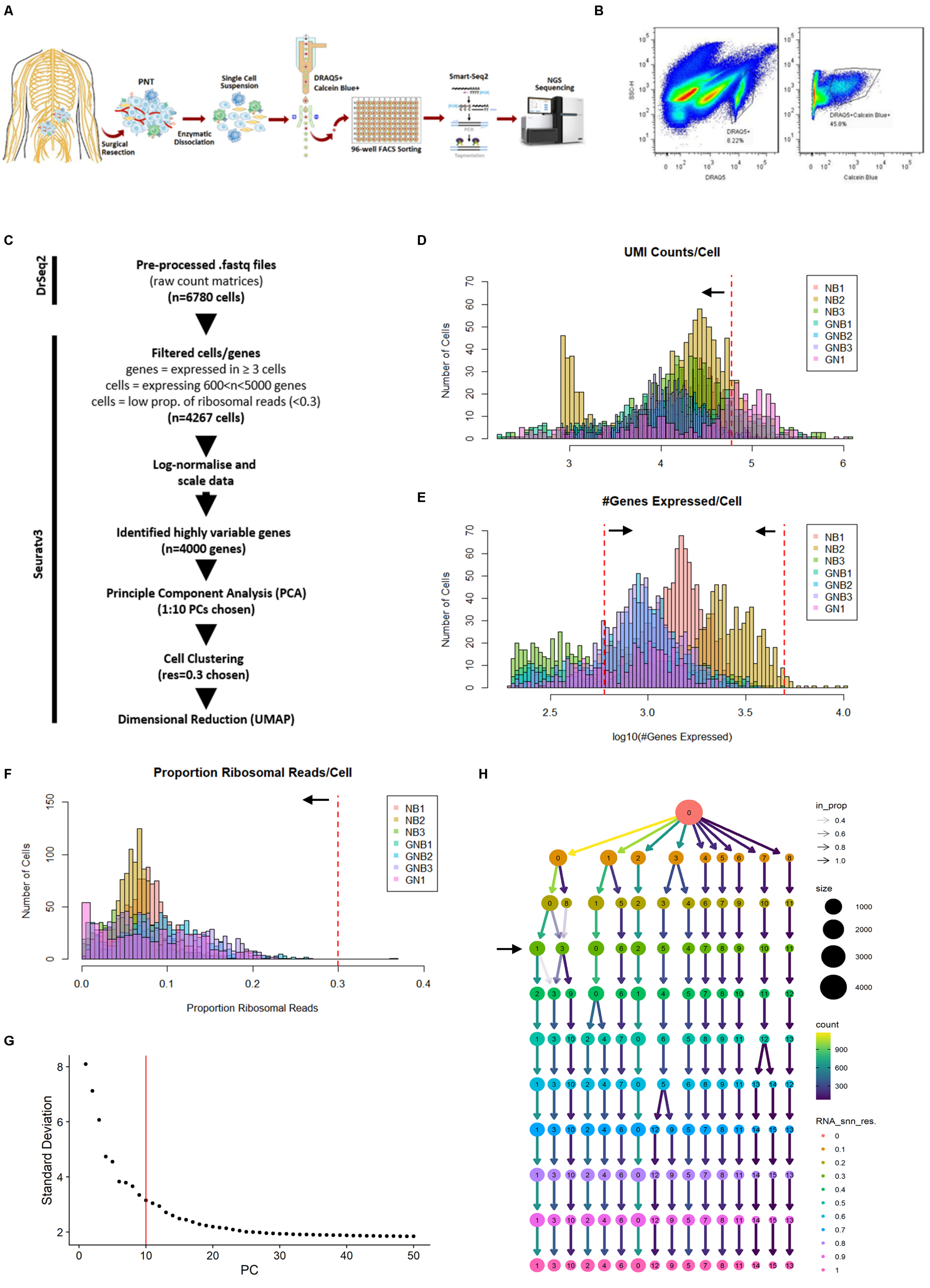

Supplementary Figure 1 (2 of 3)

I

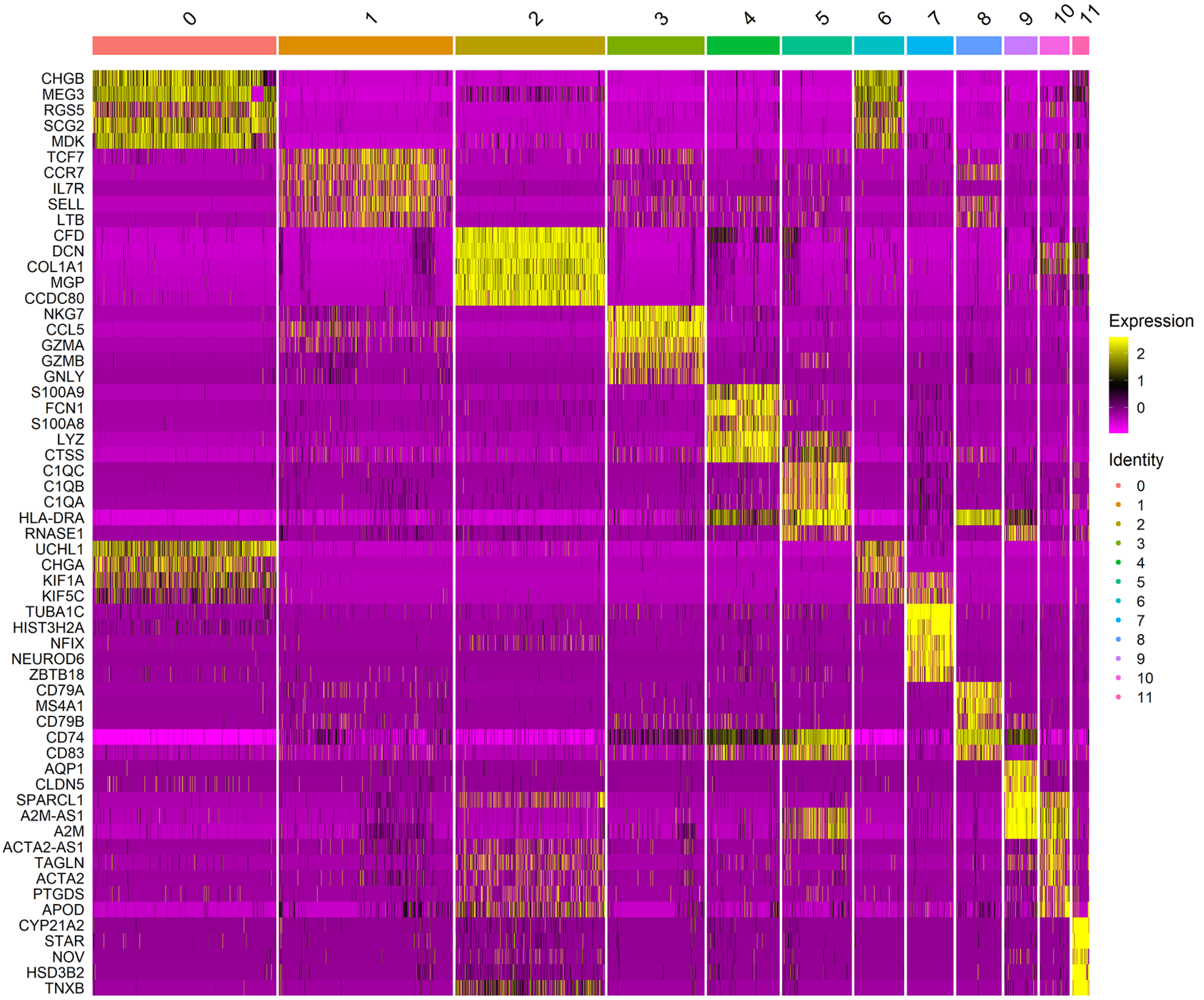


J

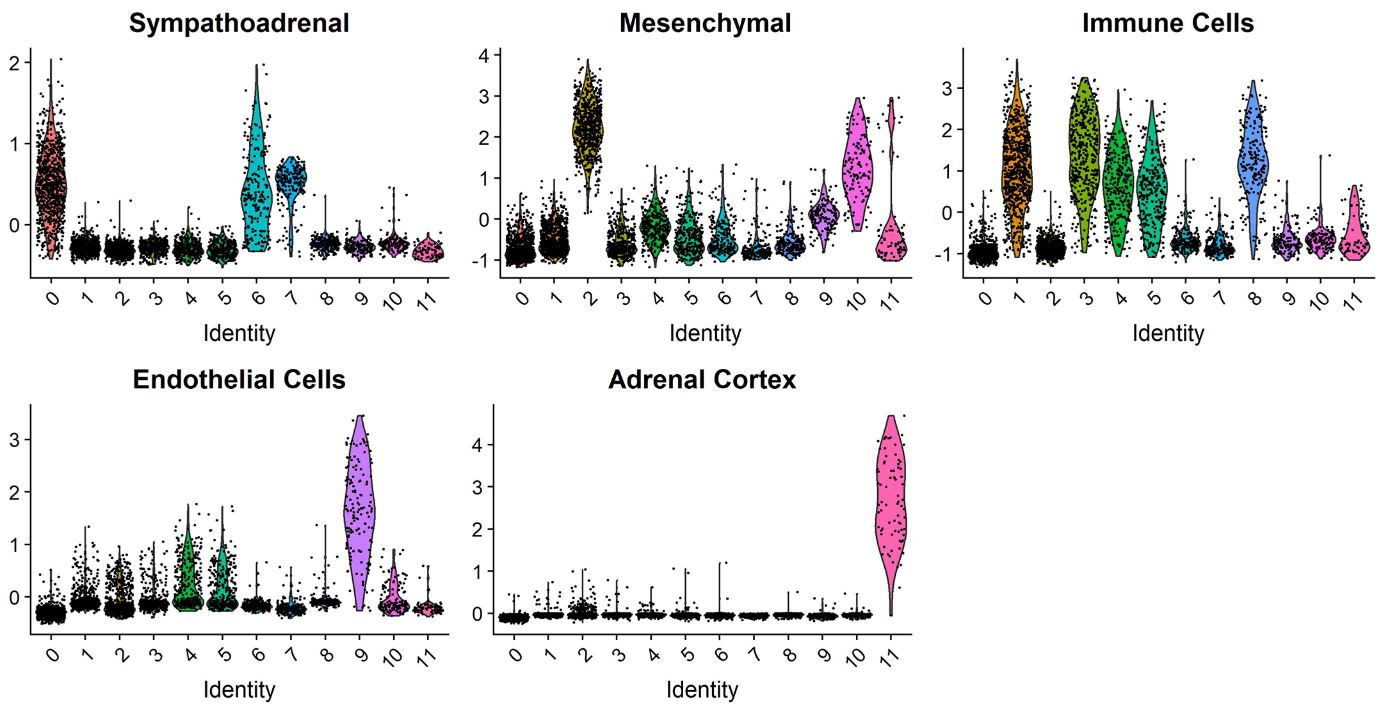

**Supplementary Figure 1 (3 of 3)**

**K**

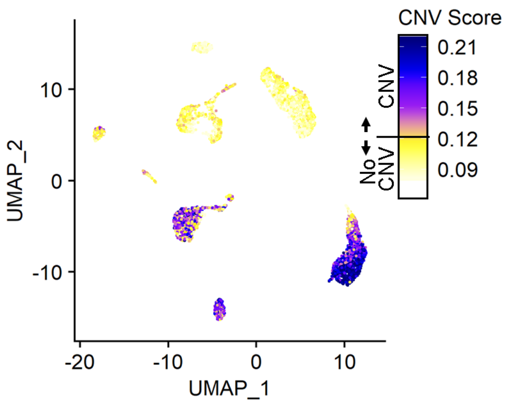


**L**

**NB1**

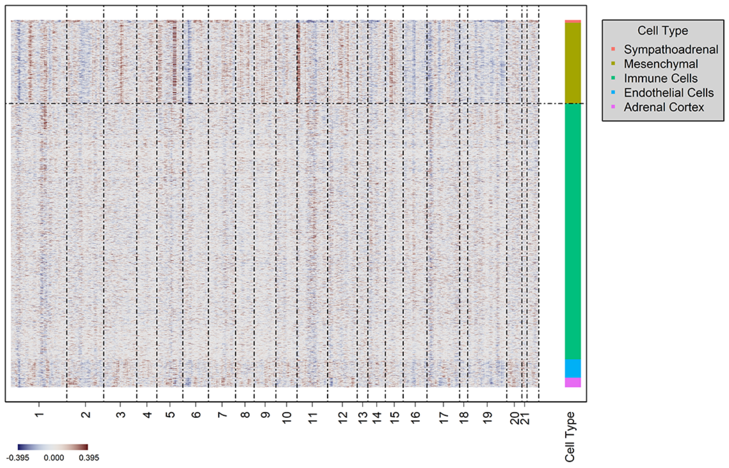


**NB3**

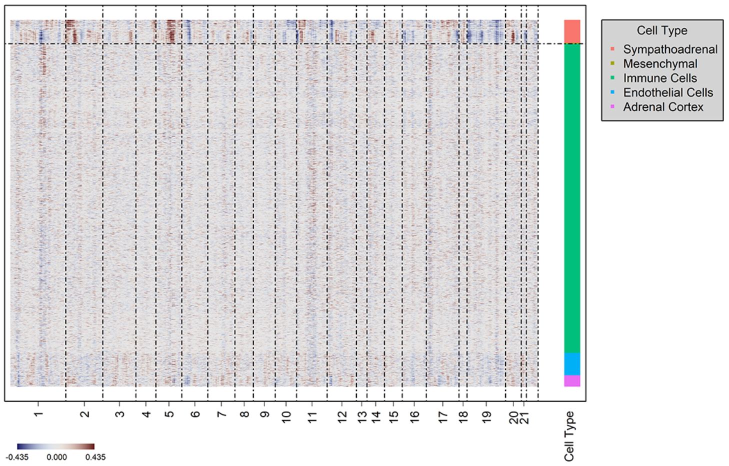


**GNB1**

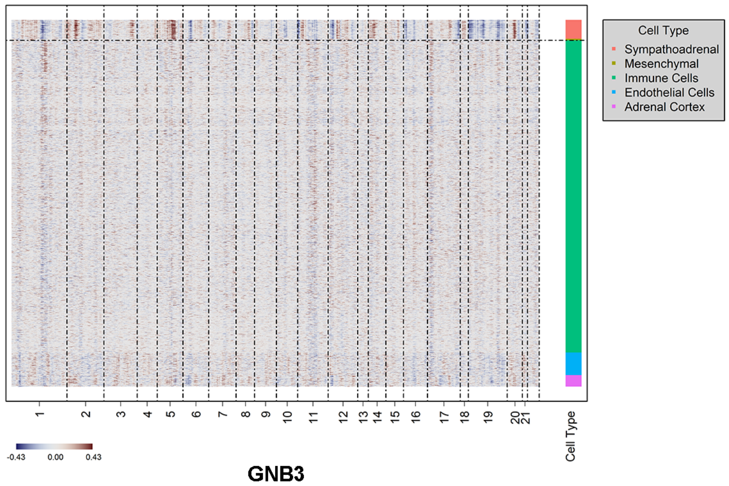


**GNB2**

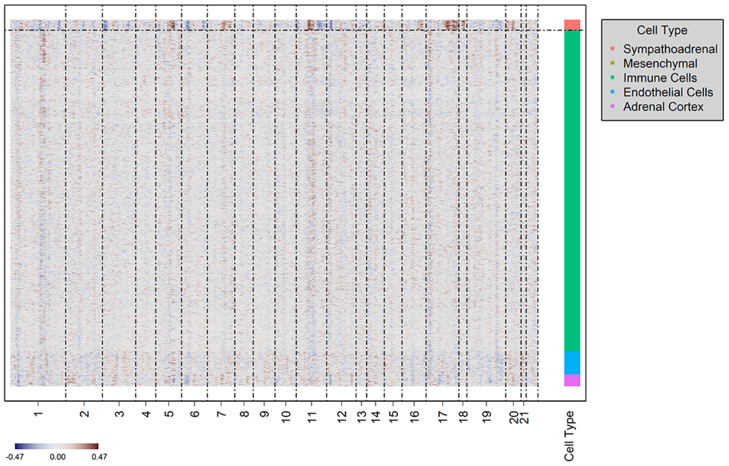


**GNB3**

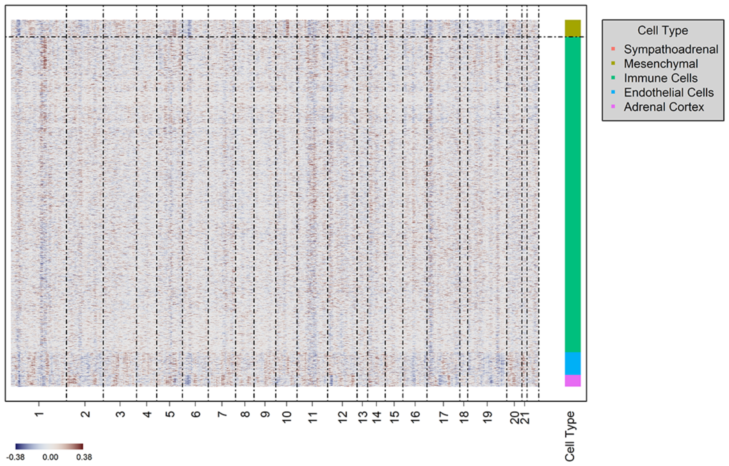


**GN1**

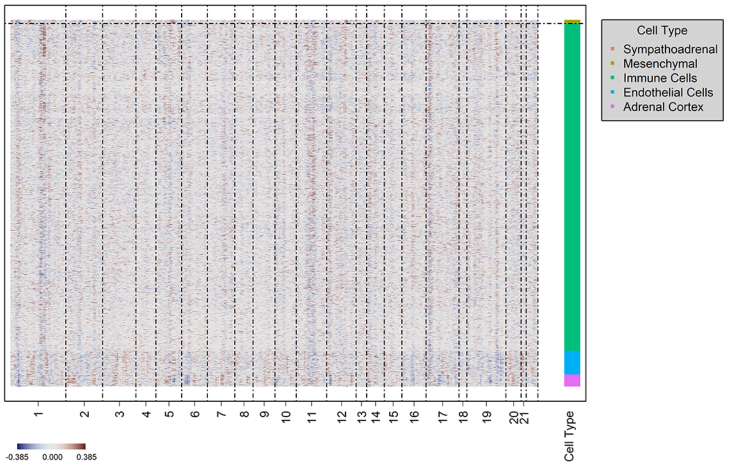

**Supplementary Figure 2 (1 of 1)**

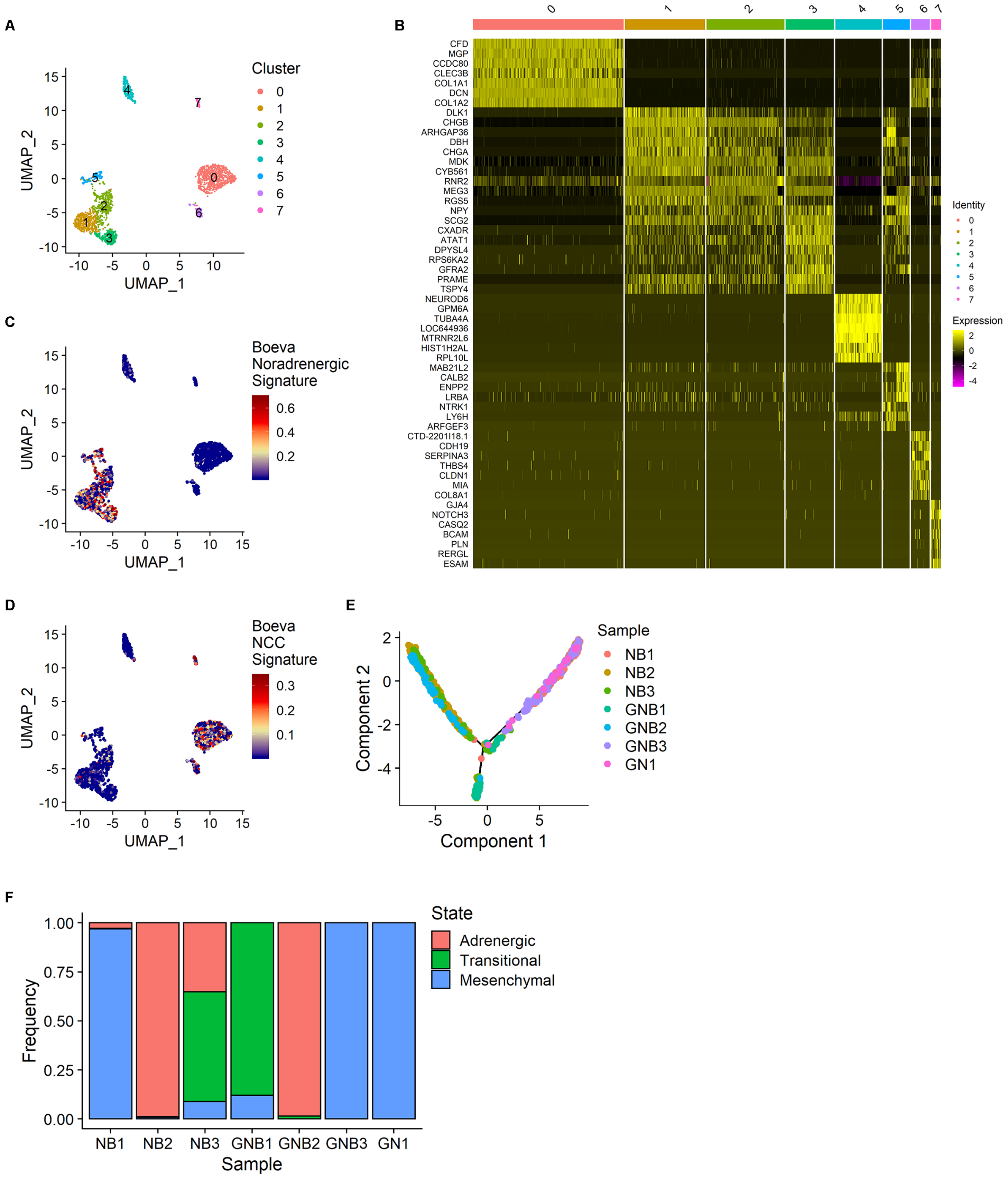

**Supplementary Figure 3 (1 of 2)**

A

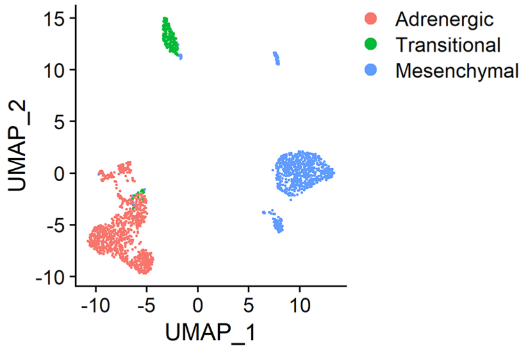


B

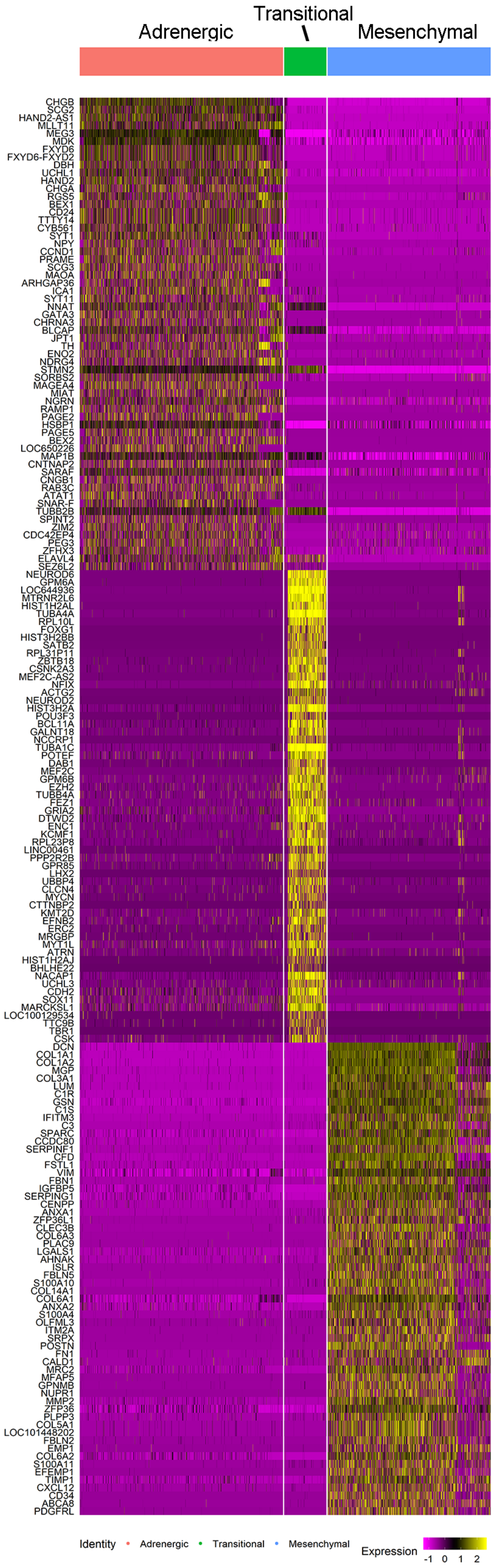


C

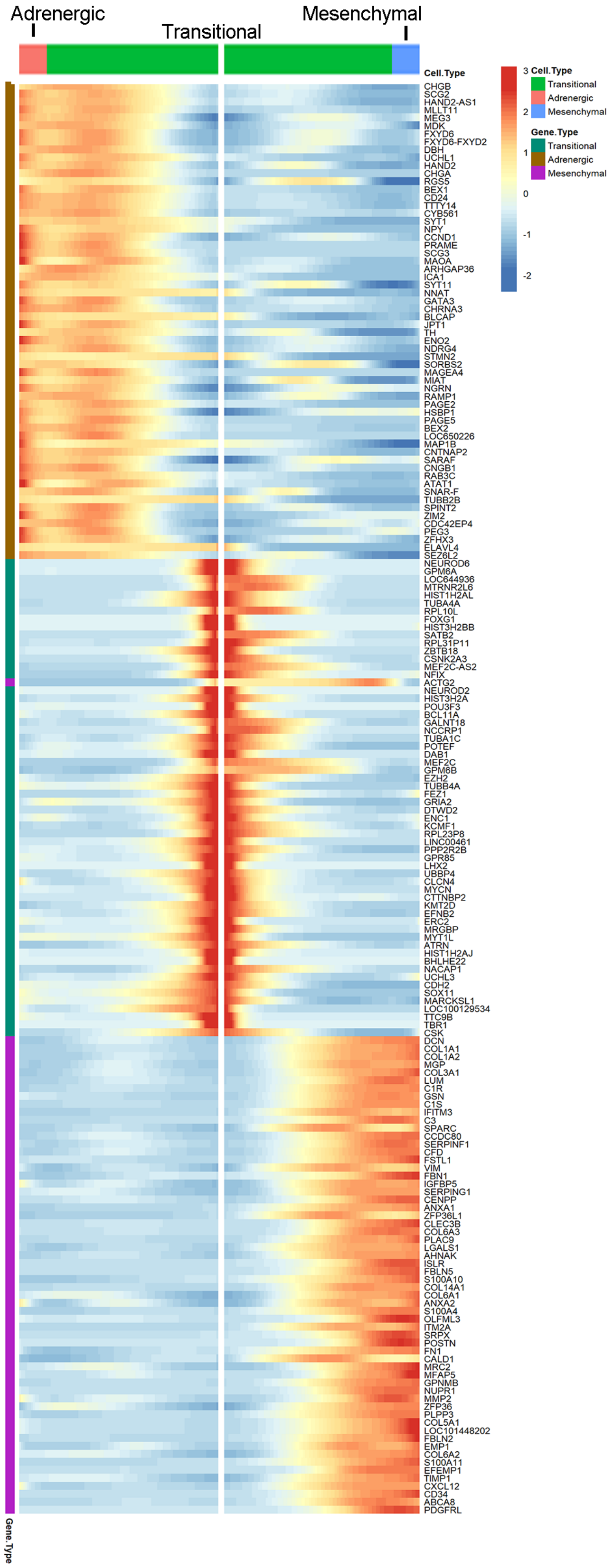

**Supplementary Figure 3 (2 of 2)**

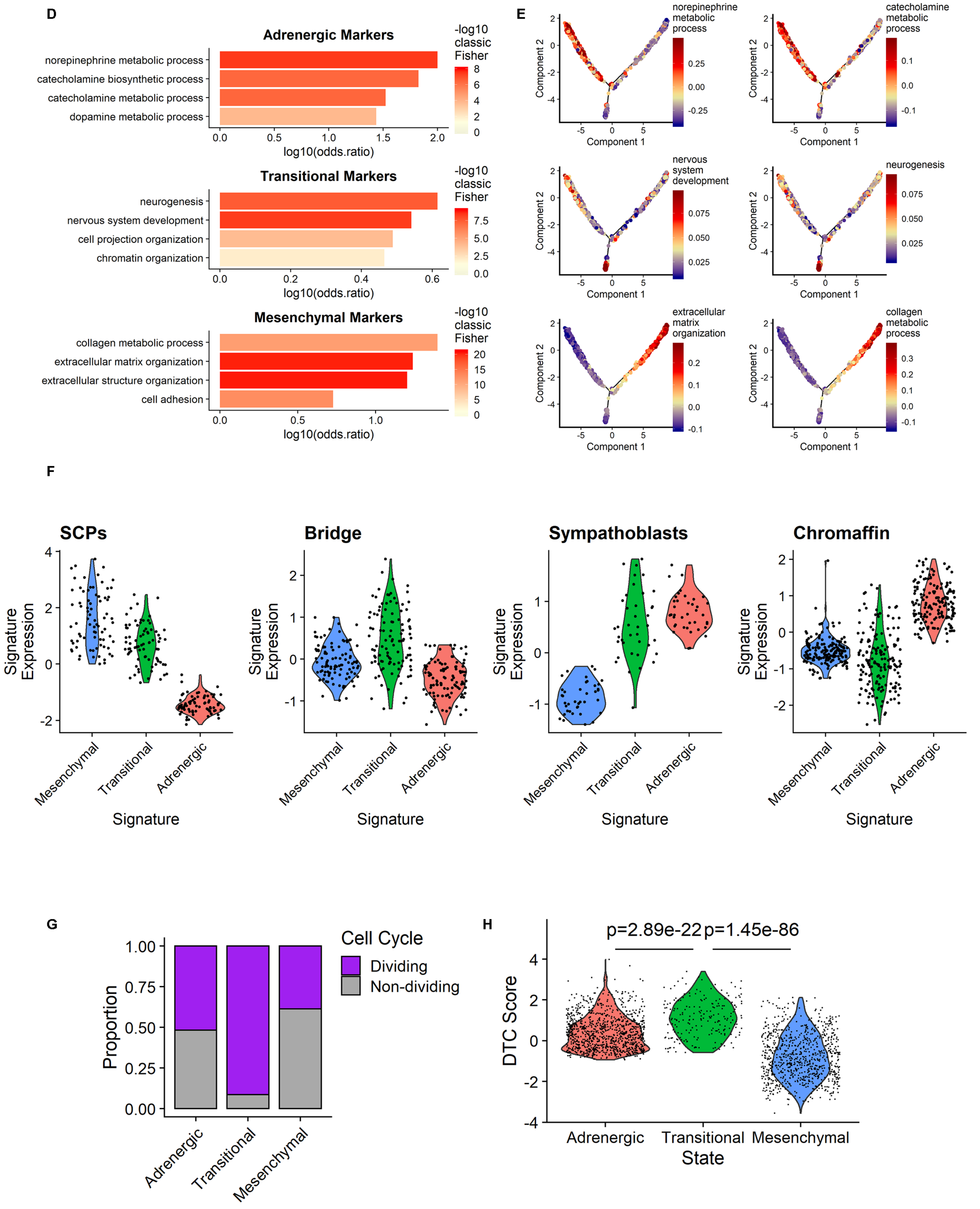

Supplementary Figure 4 (1 of 2)

A

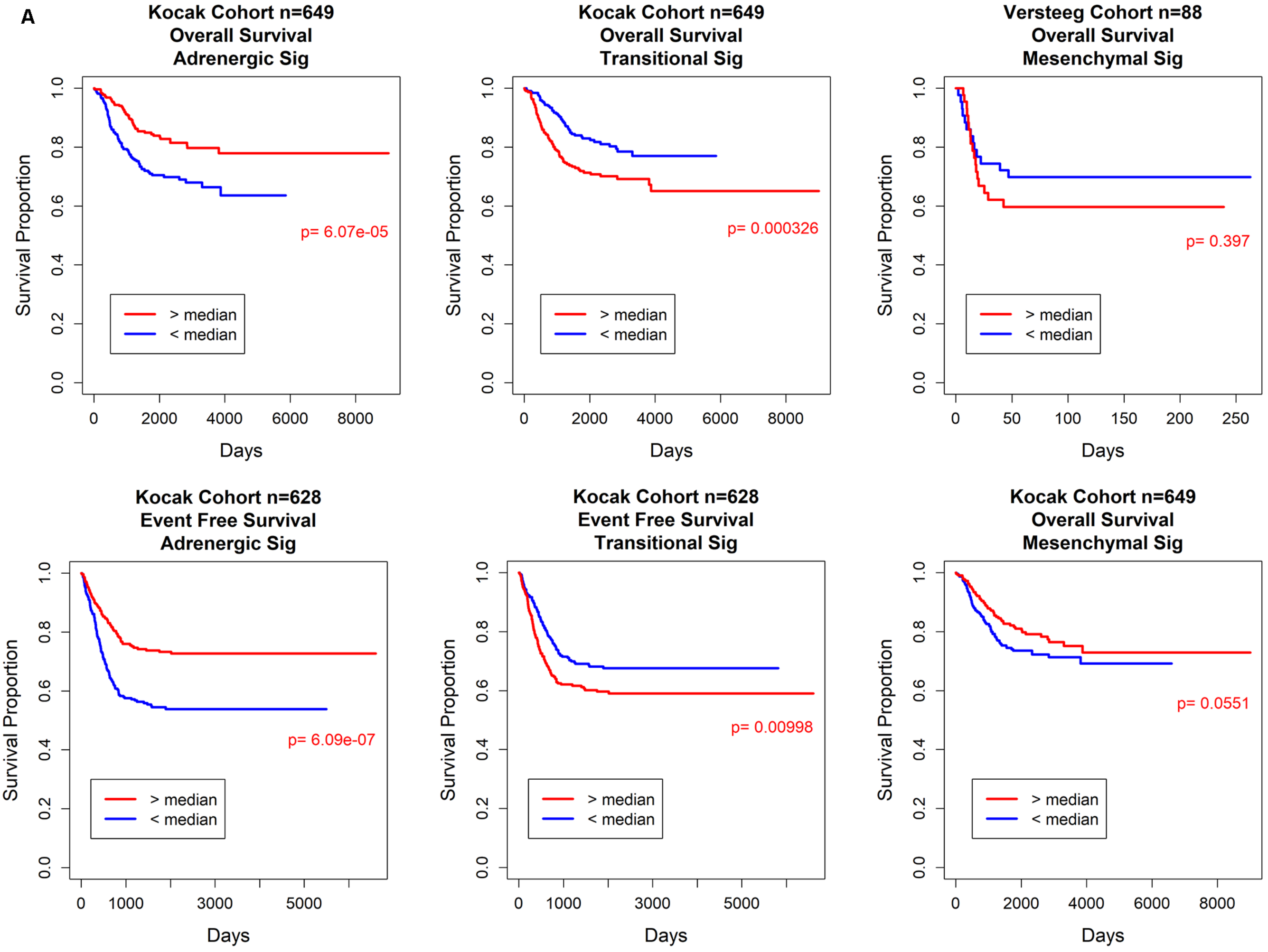


B

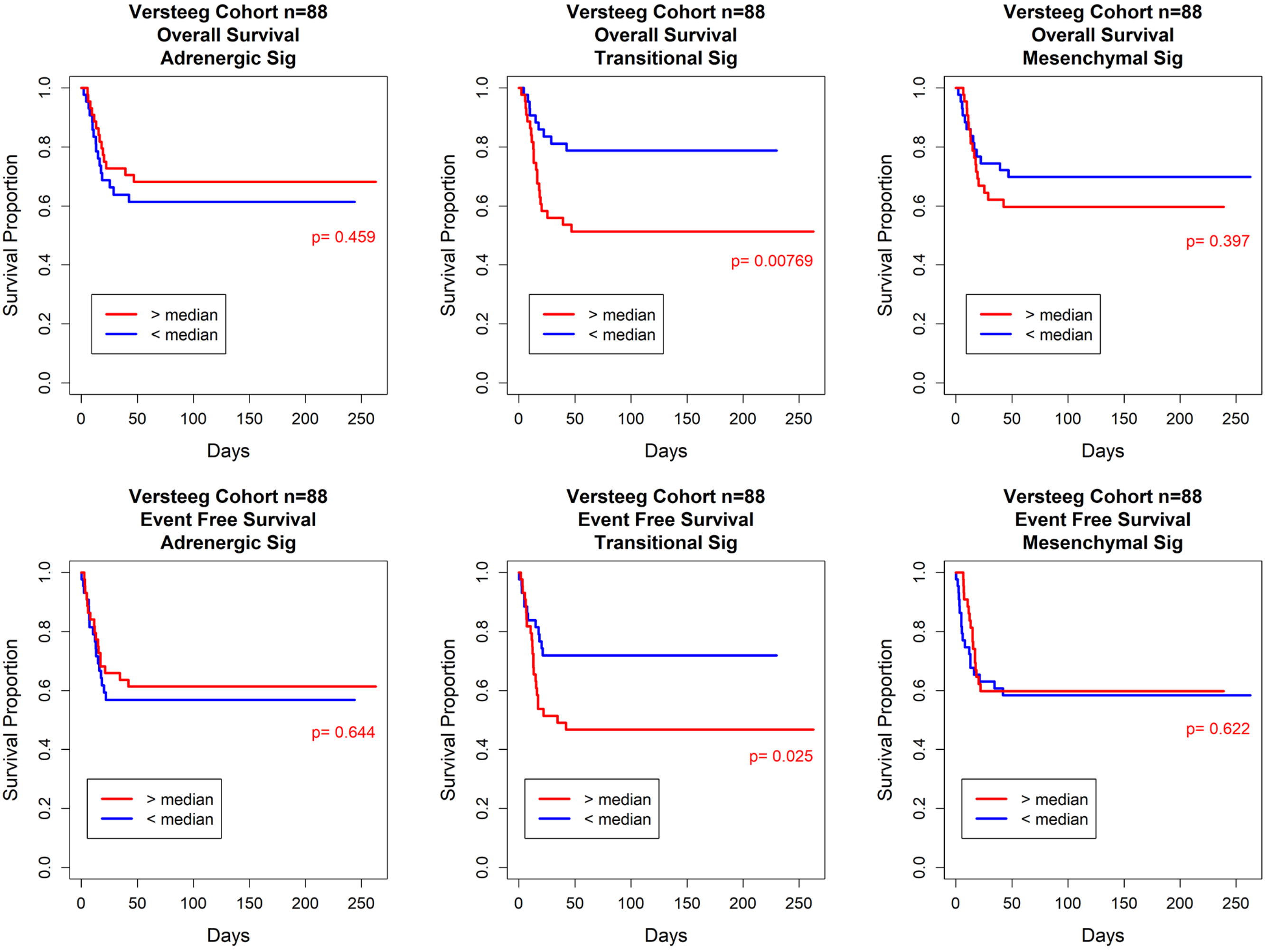

**Supplementary Figure 4 (2 of 2)**

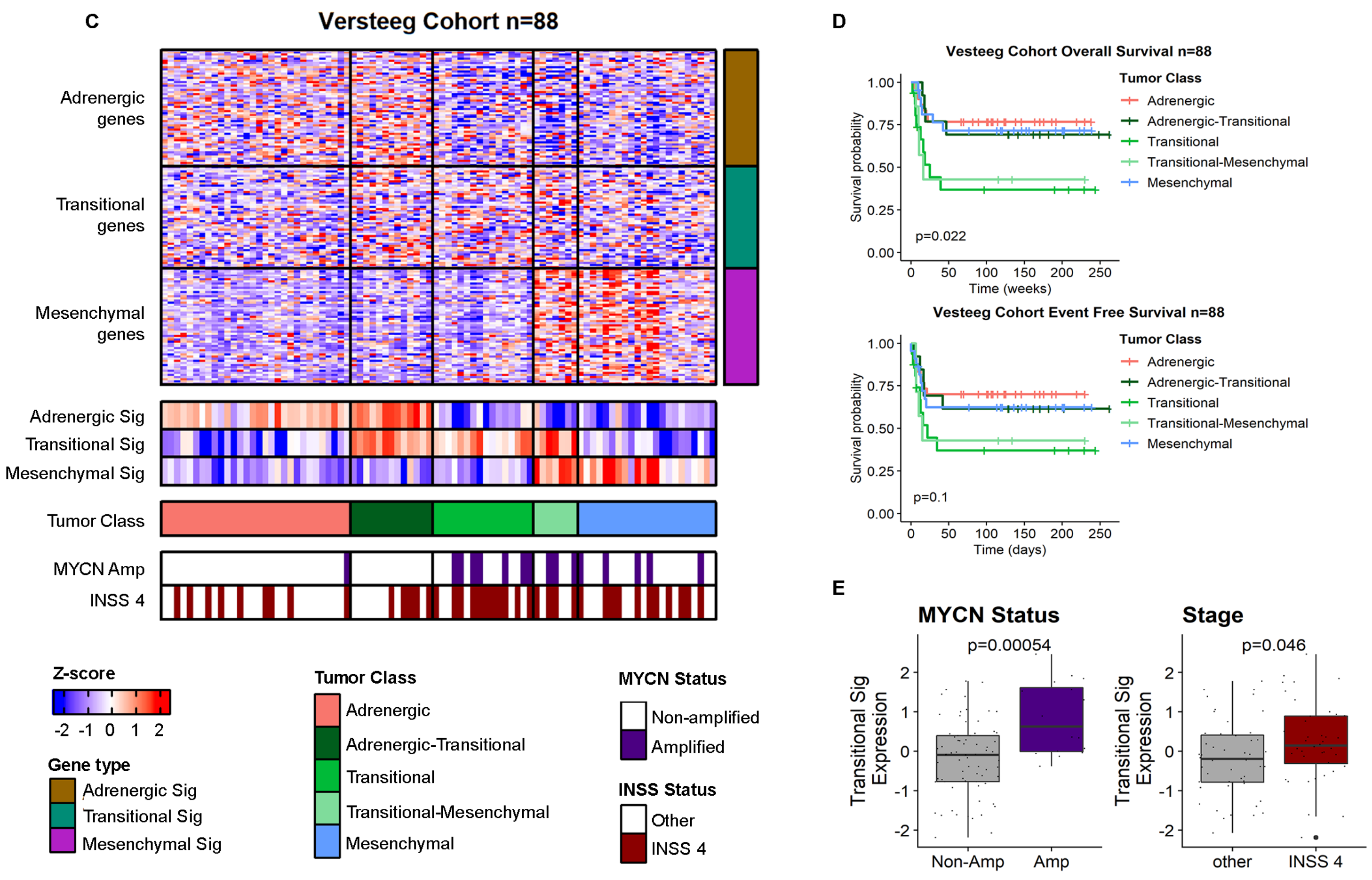

**Supplementary Figure 5 (1 of 2)**

**A**

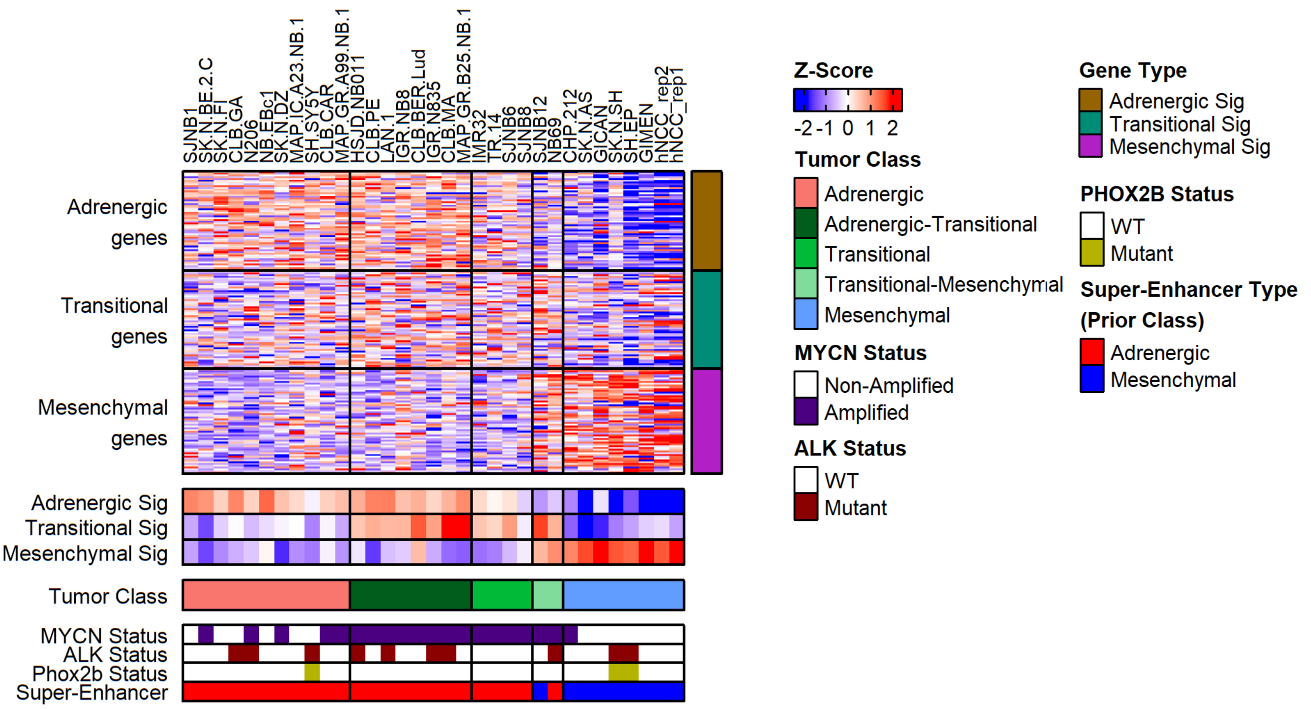


**B**

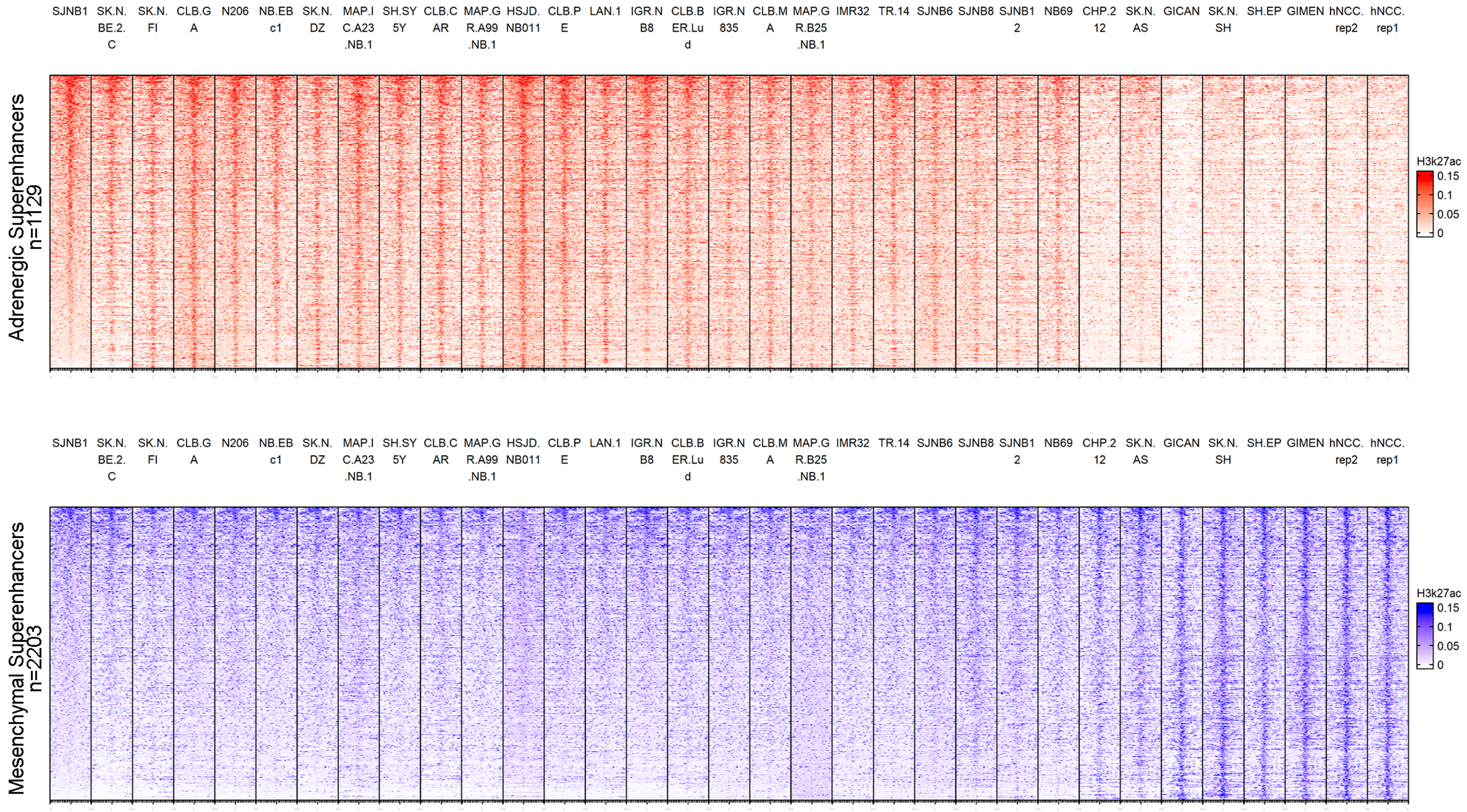


**C**

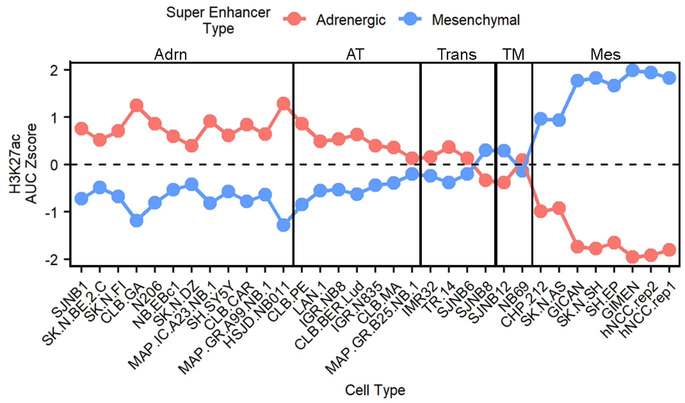

### Supplementary Figure 5 (2 of 2)

D

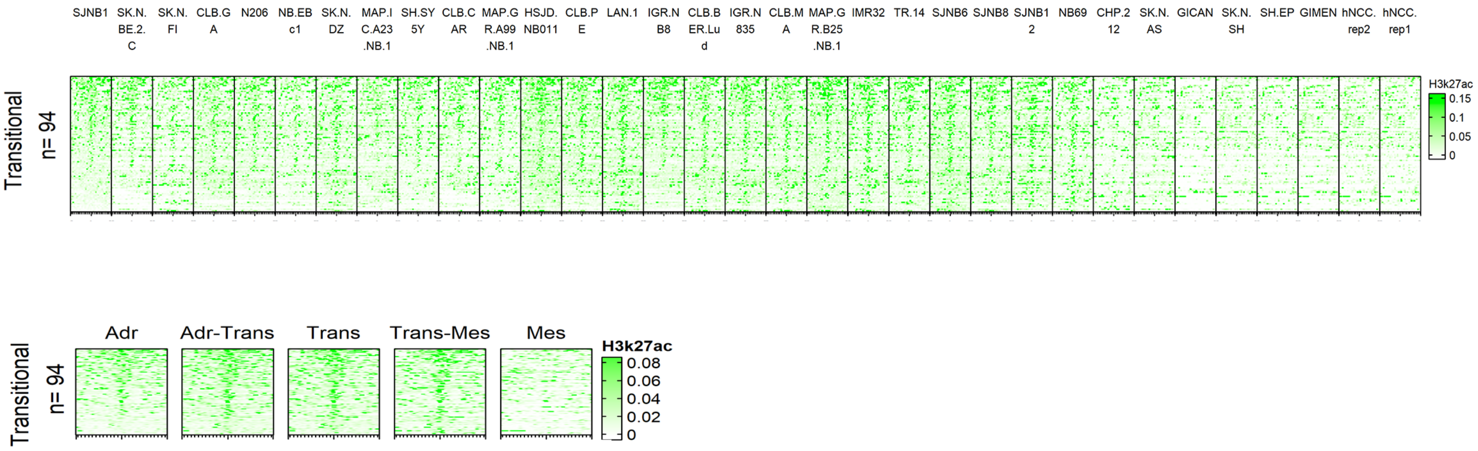


E

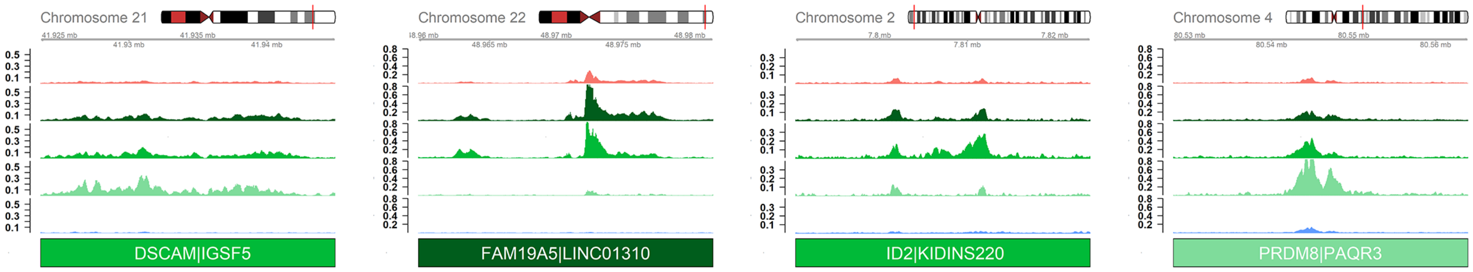

**Supplementary Figure 6 (1 of 1)**

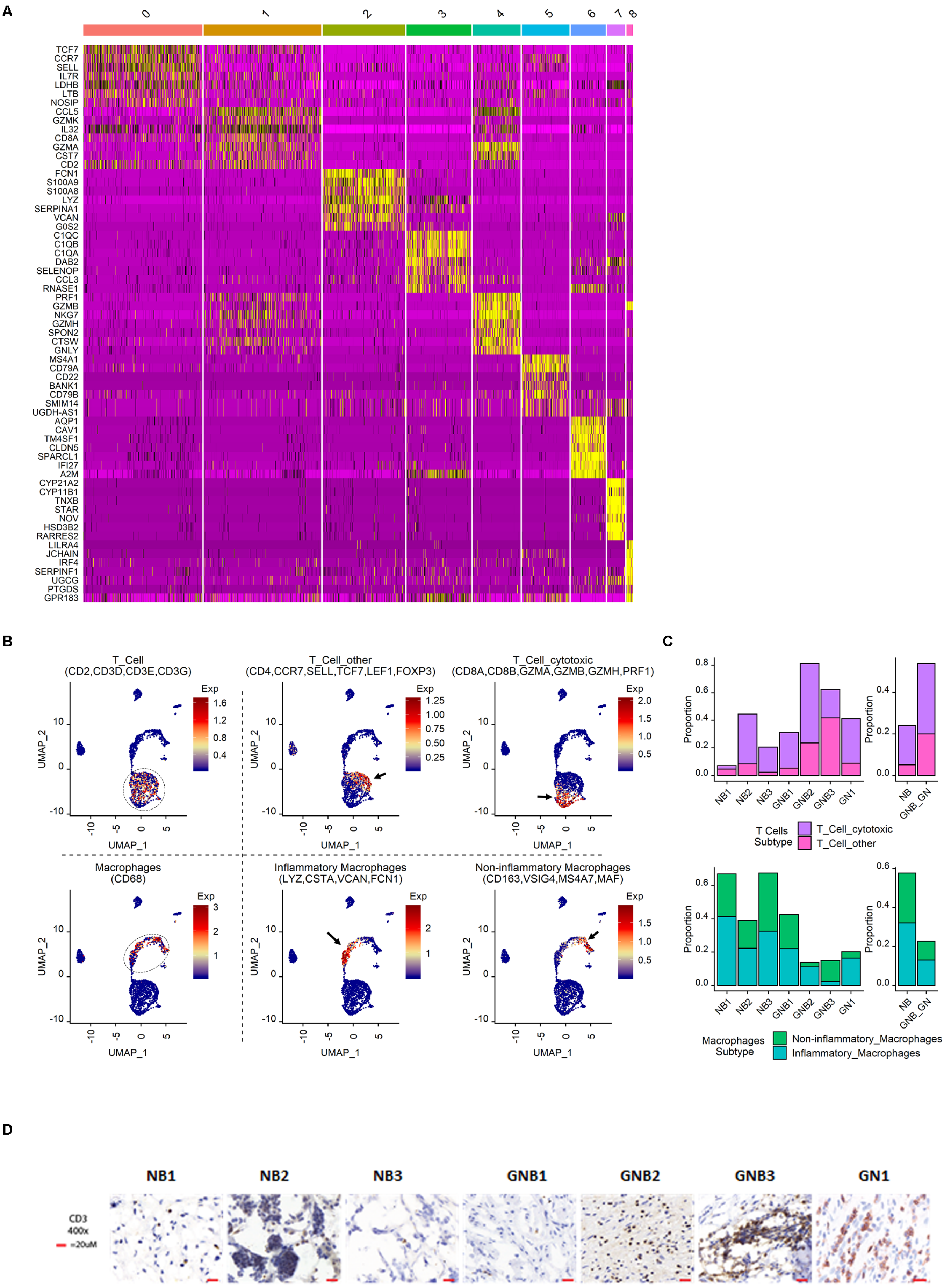

Supplementary Figure 7 (1 of 2)

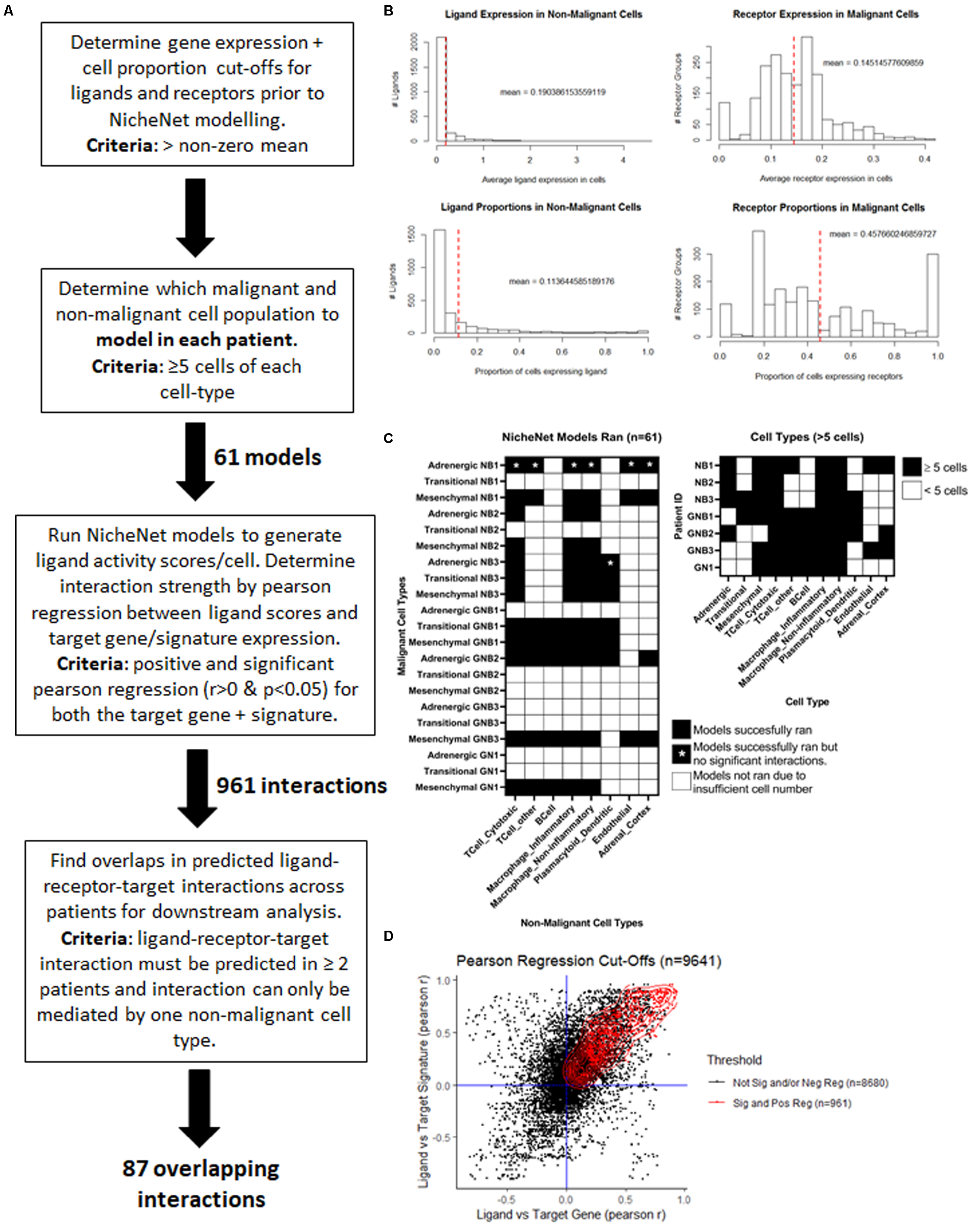

E

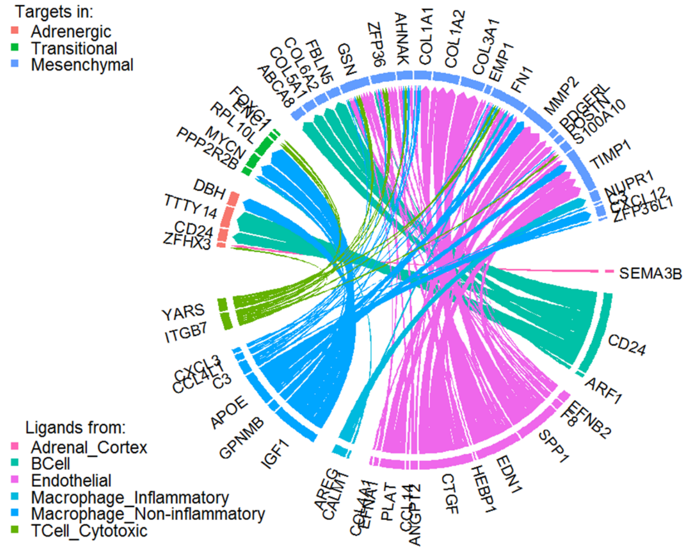
Targets in:
Adrenergic
Transitional
Mesenchymal
Ligands from:
Adrenal_Cortex
BCell
Endothelial
Macrophage_Inflammatory
Macrophage_Non-inflammatory
TCell_Cytotoxic
ZFP36
AHNAK
COL1A1
COL1A2
COL3A1
EMP1
FN1
MMP2
TIMP1
NUPR1
SEMA3B
CD24
ARF1
SPP1
EDN1
HEBP1
CTGF
PLAT
IGF1
GPNMB
APOE
C3
ITGB7
YARS
ZFHX3
CD24
TTTY14
DBH
PPP2R2B
MYCN
GSN
FBLN5
ABCA8

F

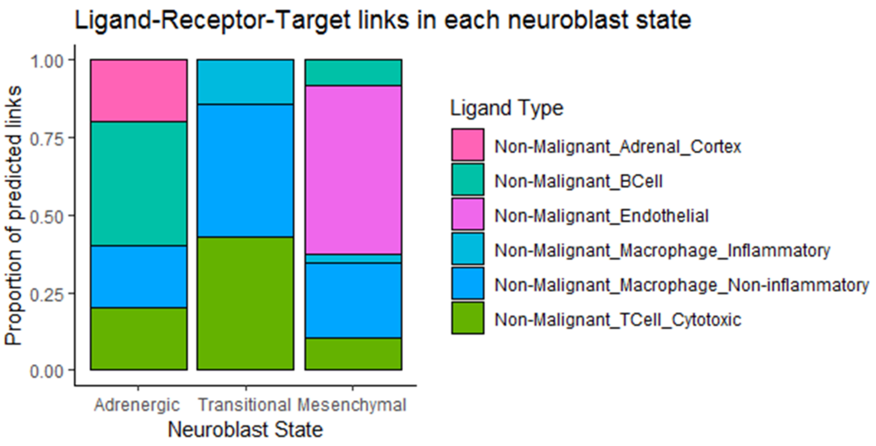
Ligand-Receptor-Target links in each neuroblast state
Proportion of predicted links
1.00
0.75
0.50
0.25
0.00
Adrenergic
Transitional
Mesenchymal
Neuroblast State
Ligand Type
Non-Malignant_Adrenal_Cortex
Non-Malignant_BCell
Non-Malignant_Endothelial
Non-Malignant_Macrophage_Inflammatory
Non-Malignant_Macrophage_Non-inflammatory
Non-Malignant_TCell_Cytotoxic

**Supplementary Figure 8 (1 of 1)**

**A**

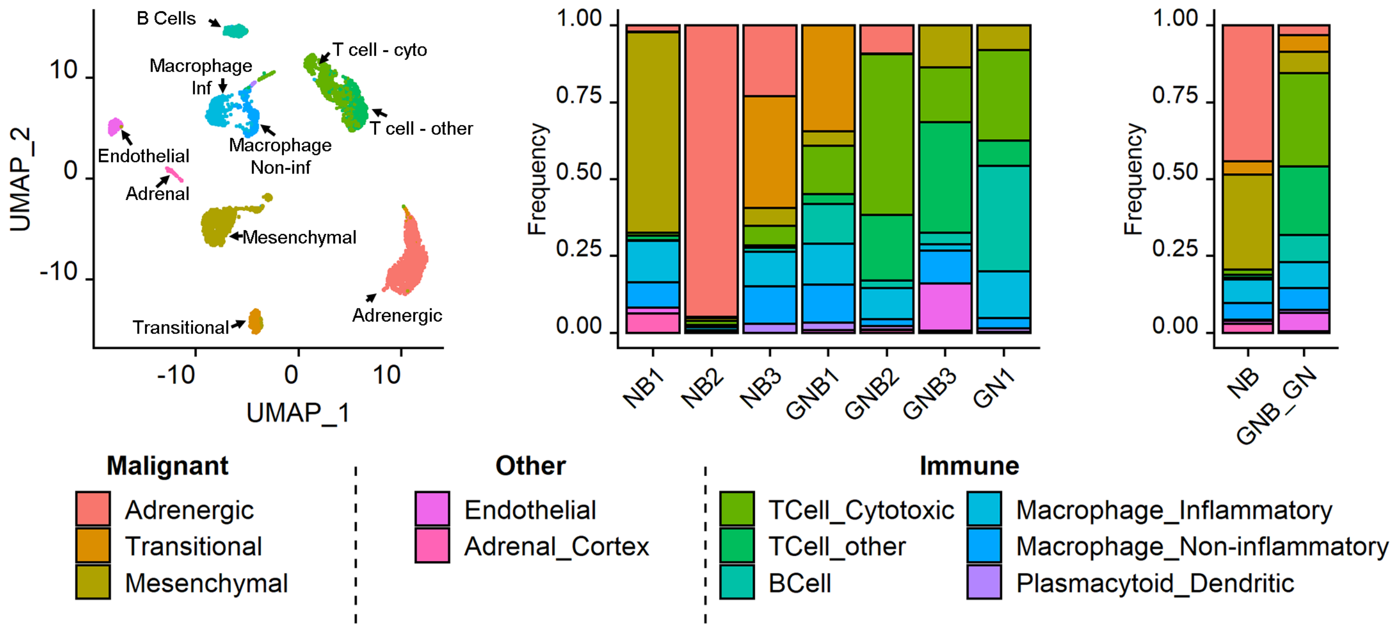


**B**

| | | NB1 | NB2 | NB3 | GNB1 | GNB2 | GNB3 | GN1 |
|---|---|---|---|---|---|---|---|---|
| **Histology** | | Neuroblastoma | Neuroblastoma | Neuroblastoma | Ganglio-Neuroblastoma: nodular | Ganglio-Neuroblastoma: nodular | Ganglio-Neuroblastoma : intermixed | Ganglioneuroma |
| **INRG Risk** | | Intermediate | High | High | High | Intermediate | NA | NA |
| **Malignant/Non-malignant ratio** | | +++ | ++++ | +++ | ++ | + | + | + |
| **Malignant Class** | | M | A | A/T | T/M | A | M | M |
| A | Adrenergic | + | +++ | ++ | - | +++ | - | - |
| T | Transitional | - | - | ++ | +++ | + | - | - |
| M | Mesenchymal | +++ | - | + | + | - | +++ | +++ |
| **Non-Malignant Class** | | M_pan | M_pan/T_pan | M_pan | T_pan/M_pan | T_pan | T_pan | T_pan |
| TC | T-cytotoxic | - | ++ | + | ++ | +++ | ++ | ++ |
| TO | T-other | - | ++ | - | ++ | +++ | +++ | ++ |
| IM | Inflammatory-macrophage | +++ | ++ | +++ | ++ | + | - | + |
| NIM | Non-Inflammatory-macrophage | +++ | ++ | +++ | ++ | - | + | - |
